# Characterizing the dual regulation of *Neisseria gonorrhoeae tdfJ* in response to zinc and iron

**DOI:** 10.1101/2024.08.01.606247

**Authors:** Sandhya. Padmanabhan, Julie Lynn Stoudenmire, Alexis Hope Branch, Cynthia Nau Cornelissen

## Abstract

Gonorrhea is a sexually transmitted infection, caused by the bacterial pathogen *Neisseria gonorrhea* (*Ngo*) and affects millions of individuals of all age groups across the globe every year. Infection with *Ngo* does not result in protection and no effective vaccine has been developed, leaving antibiotics as the only treatment option. With the emergence of strains showing high levels of antibiotic resistance, there is an urgent need for development of novel therapeutics for disease prevention. During pathogenesis the host employs nutritional immunity, to restrict important transition metals such as zinc away from *Ngo*. This process is counteracted in *Ngo* by the production of highly efficient zinc import TonB-dependent Transporters (TDTs) which are promising vaccine antigens and zinc shuttle ABC transporters found to be important for intracellular survival. In *Ngo* zinc homeostasis and transport proteins are regulated by the Zinc uptake regulator (Zur) which represses transcription in the presence of zinc and activates transcription in the absence of zinc. In this study, characterize the promoter elements of the zinc import TDT, *tdfJ*, which results in dual regulation by zinc and iron. We characterize specificity and binding affinities for regulation of *tdfJ* by a second regulator, Ferric uptake regulator (Fur) in response to iron. The response of *tdfJ* to both iron and zinc and its potential to be an important invasin, makes it an attractive candidate to investigate female genital tract infections. The female genital tract is a conglomerate of these conditions and infections here are often asymptomatic. Taken together, this research provides important knowledge on the regulation of virulence mechanisms in response to zinc, which will aid in the development of therapeutics and an efficacious vaccine against a gonococcal infection.

## INTRODUCTION

*Neisseria gonorrhoeae* (*Ngo*) causes the sexually transmitted infection gonorrhea, that affects predominately younger adults, both men and women, worldwide. *Ngo* is a major public health concern and the World Health Organization (WHO) estimates that there were 82.4 million new cases as of the year 2020, a 118% increase since 2007 (1). There is no protective immunity, no effective vaccine, and historically only antibiotics have been used to treat infection. The current recommended treatment is a third-generation cephalosporin, ceftriaxone, due to the increasing numbers of resistant strains to all the other classes of antibiotics used over the decades (2). However, with the threat of untreatable *Ngo*, the Center for Disease Control and Prevention (CDC) specified *Ngo* a superbug and an urgent threat pathogen (3). This is alarming and there is an urgent need for vaccine and therapeutics to treat gonorrhea. Gonococcal infection in men, primarily results in an inflammatory response that is accompanied by a purulent discharge with polymorphonuclear leukocytes (PMNs), which leads to urethritis (4, 5). In women, gonococcal infection primarily results in cervicitis, but is often asymptomatic or accompanied by nonspecific symptoms (6). As a result of asymptomatic infection, untreated infection can cause serious irreversible sequelae such as pelvic inflammatory disease (PID), ectopic pregnancies, and even infertility (5, 7). This molecular difference in pathogenesis and immune response to infection in men and women is not well understood and emphasizes a gap in knowledge on the mechanisms by which *Ngo* adapt to these two distinct environments.

The human host employs strategies for immune clearance of *Ngo* including a robust inflammatory response with an influx of neutrophils or PMNs that entrap pathogens in neutrophil NETs (5, 8). The gonococcus has, however, been shown to be escape the harsh neutrophil NETs and in turn evade capture and phagocytosis. In fact, intact *Ngo* have been noted in these purulent urethral or cervical exudates from patients presenting with gonorrhea. These host immune cells also produce antimicrobial agents and proteins to limit essential nutrients for bacterial survival by a process termed nutritional immunity (9). *Ngo* has developed highly efficient transport systems, called TonB-dependent transporters (TDTs), to overcome this nutrient limitation and scavenge for metals by metal piracy from host proteins, wreaking havoc in the host. There are 8 known TDTs in *Ngo* of which 2 are characterized as zinc importers. Zinc import and homeostasis is a key feature in maintaining virulence and pathogenesis in several pathogens, including the gonococcus. The two, zinc import TDTs are TdfH and TdfJ, which function with an inner membrane ZnuABC transporter to cooperatively import zinc. TdfH binds S100A8/A9 (calprotectin), which is enriched inside human neutrophils to hijack zinc, while TdfJ binds S100A7 (psoriasin) secreted by mucosal epithelia, found inside the female cervical and vaginal mucosa (10–13). These TDTs are conserved across all strains of *Ngo* and have been implicated as important vaccine antigens (14). ZnuA has also been demonstrated to be essential for gonococcal survival in an epithelial infection model (11), suggesting zinc import is crucial for survival. At the molecular level these zinc import proteins are regulated by the Zinc uptake regulator (Zur) which senses internal and external cellular zinc levels in order to maintain zinc homeostasis and overcome zinc starvation or toxicity imposed by the host (15). When faced with zinc toxicity, Zur binds the excess Zn^2+^ ions and blocks transcription of Zur regulated genes including *tdfH*, *tdfJ* and *znuABC* (16, 17). During zinc-starvation, Zur does not repress these promoters and transcription, and protein production is enhanced to combat the zinc stress. Zur carefully modulates the expression of these genes by binding to specific regions on their promoters, which is the subject of this study.

During pathogenesis, *Ngo* face different levels of metal environments, including zinc-dependent toxicity and starvation depending on the host sex and tissue type (5, 18, 19). The female genital tract is more zinc starved than the male genital tract (20). Given the fact that women tend to have a higher percentage of asymptomatic infections, coupled with the variation in zinc levels between the male and female host, it is important to ask the question if *Ngo* utilize one TDT over the other, and if this impacts disease progression between different hosts. In this study, we sought to characterize the extent to which these virulence genes respond to zinc and are regulated by Zur along with the promoter elements that modulate their expression levels. We identify a candidate gene that has a dual regulatory mechanism and speculate its possible role in gonococcal infections in women.

## RESULTS

### *tdfJ* and *znuA* Show Significant Changes in Gene Expression by RT-qPCR

We previously demonstrated that the gonococcal zinc import TDTs, TdfH and TdfJ and their cognate ABC transporter, ZnuABC, are all regulated at the translational level by Zur, in a zinc-dependent manner (13). However, the extent of transcriptional regulation by Zur on these genes under excess and low zinc conditions has not been studied. Here we sought to quantify the level of regulation by Zur on these *tdfJ* and *tdfH* by RT-qPCR. We grew the FA1090 (WT) and an isogenic *zur* mutant (*zur*) in chelex-treated defined medium (CDM), under excess zinc (20 µM Zn_2_SO_4_) or low zinc (7 µM TPEN) conditions. TPEN was added as a zinc-specific chelator to induce zinc-starvation. Cells were collected during exponential phase and mRNA and whole cell lysates were processed for gene expression and protein analysis. RT-qPCR was performed to measure the log_2_-fold-change for gene expression for *tdfH*, *tdfJ*, and *znuA*. Gene expression levels for *ngo1049* and *tbpB* were also measured as a positive and negative control, respectively. RT-qPCR expression for these genes were normalized to *rmpM* expression levels, a house keeping gene not regulated by zinc or Zur. *tdfJ* expression was 24-fold higher in the absence of zinc in WT cells as compared to the *zur* mutant (Figure 1b). *tdfJ* expression in the *zur* mutant under excess zinc is comparable to the expression in the absence of zinc, indicating a Zur-dependent mechanism of regulation. *tdfH* expression is increased 3-fold in the absence of zinc in WT cells, and periplasmic zinc binding protein, *znuA*, was increased 38-fold, with no significant difference in the *zur* mutant (Figure 1a and 1d, respectively). *znuA* is necessary for zinc assimilation during attachment of *Ngo* (11), making its tight regulation by Zur important during infection. In WT cells, *ngo1049* expression was increased 46-fold in the absence of zinc, which was not seen in the *zur* mutant (Figure 1c). *tbpB*, however, showed no significant difference in the WT or *zur* mutant (Figure 1e). Western blotting for whole cell lysates showed complementary protein expression levels to RT-qPCR. TdfJ was only produced in the absence of zinc in the wildtype while having no qualitative difference in protein levels in the *zur* mutant in the presence or absence of zinc (Figure 1a). TdfH did not appear to be fully repressed in the presence of zinc in the wildtype, which was shown in a previous study to be a result of strain-specific difference between FA1090 and FA19 *Ngo* (Figure 1b) (13). There was no difference in TdfH protein levels in the *zur* mutant in the presence or absence of zinc (Figure 1b). ZnuA protein levels were similarly expressed to complement the RT-qPCR expression data. ZnuA was produces in the absence of zinc in the wildtype with no difference in the zur mutant in protein levels for zinc abundance or zinc depletion (Figure 1c). One thing to note is the small yet-significant difference in *tdfJ* and *znuA* expression in the *zur* mutant between high and low zinc levels (Figure 1). This difference is however not visible in their protein expression levels in the *zur* mutant, suggesting another layer to their transcriptional regulation. This difference in the *zur* mutant was also seen in the positive control *ngo1049* expression in the *zur* mutant which was further recapitulated in the western data with a visible difference in the *zur* mutant Ngo1049 production between zinc-excess and low zinc conditions (Figure 1d). The transcriptional and translational regulation by Zur, overall are complementary, however suggesting a stricter transcriptional layer in regulation by Zur in response to zinc.

**Figure 1.**
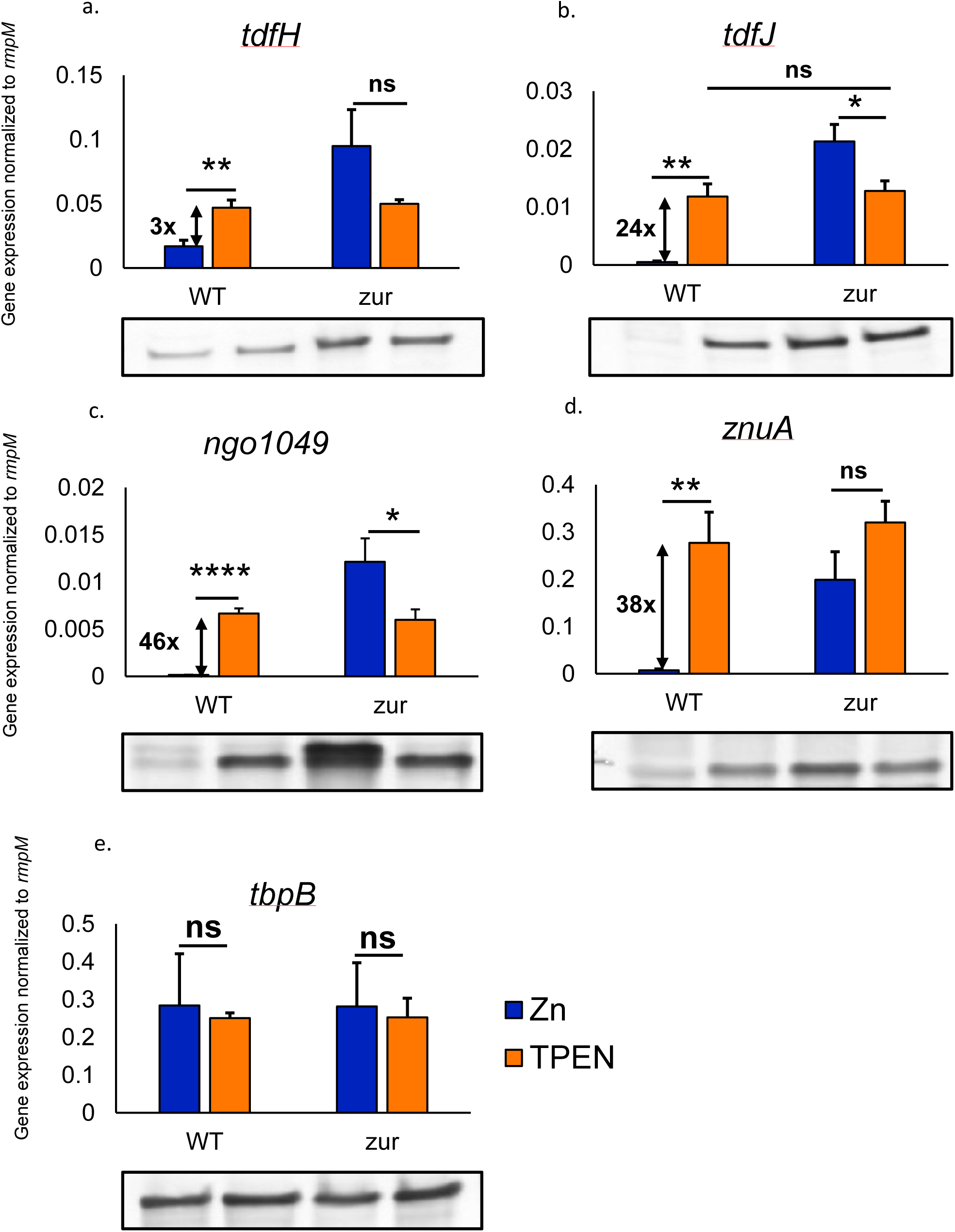
*tdfH* (a), *tdfJ* (b), *ngo1049* (c) and *znuA* (d) are all zinc-regulated by Zur. Gene expression was normalized to *rmpM* before plotting bar graphs. *tbpB* (e) expression was also computed as a negative control for zinc-regulated expression. Excess zinc condition is depicted in blue while the zinc-restricted condition in the presence of TPEN is shown in orange. 2ΔCq values from three individual biological replicates were averaged. Student’s t test was used to calculate statistical significance. The corresponding western blots for protein expression of TdfH, TdfJ, NGO1049, ZnuA and TbpB are shown below the respective RT-qPCR plots. * p<0.05,**p<0.005, **** p<0.00005, ns-not significant, p>0.05.

### Zur Binds a Consensus Region on the tdfJ Promoter and Tightly Regulates its Expression

Previously, a putative consensus sequence for Zur binding in *Neisseria meningitidis* (*Nme*) was generated (16). Using the *Nme* predicted sequence in combination with RNA Seq promoter prediction for zinc- and Zur-regulated genes an *Ngo* Zur consensus logo was generated using WebLogo software (21) (Figure 2a). In silico promoter analysis of the putative promoters of *tdfH*, *tdfJ*, *znuA*, and *ngo1049*, identified a perfect consensus Ngo Zur box in the *tdfJ* and *ngo1049* promoters (Figure 2b). The predicted Zur-box upstream of *tdfH* has a 16 % match to the *Ngo* consensus sequence and demonstrated less difference in expression between the zinc replete and zinc deplete conditions. *znuA* showed a 92 % match for the consensus Zur-box sequence and a much higher difference between zinc replete and zinc deplete conditions. Since *tdfJ* exhibited both a bigger difference in expression between the zinc replete and zinc depleted condition and a 100 % match to the Zur box consensus we further characterized its promoter elements using BPROM and PRODORIC online webtools (22, 23) (Figure 3). Interestingly, the prediction software identified two potential translational starts with an ATG encoding for a methionine within 15 base pairs of each other. There was no canonical Shine Delgarno-Ribosome Binding Site (RBS)-like sequence (AGGAGG) in front of the first ATG, however, six nucleotides upstream of the second ATG was a predicted RBS (GAGAAG) (Figure 3). To potentially identify which translational start codon is being used, we determined the transcriptional start site (+1) for the *tdfJ* promoter. We used 5’ Rapid Amplification of cDNA Ends (5’ RACE) (24) to identify the transcriptional start site for *tdfJ* using mRNA isolated from the WT + TPEN cells or the *zur* mutant + Zn cells, to maximize the *tdfJ* transcripts. Using gene specific *tdfJ* primers dC-tailed *tdfJ* cDNA was amplified and TOPO cloned into DH5α competent cells. The location of the +1 was determined as the first nucleotide preceding which the dC tailed ends from sequencing of transformed colonies. Multiple sequences from three different biological replicates (either FA1090 WT +TPEN or FA1090 *zur* + Zn 5’ RACE products) were aligned using ApE (25) and the +1 was identified as the adenosine, 25 base pairs upstream of the second ATG and 7 base pairs downstream from the predicted -10 region (Figure 3). The identified +1 is only 7 base pairs upstream of the first potential ATG (without room for an RBS upstream), therefore, the second ATG was designated as the translational start for TdfJ.

**Figure 2.**
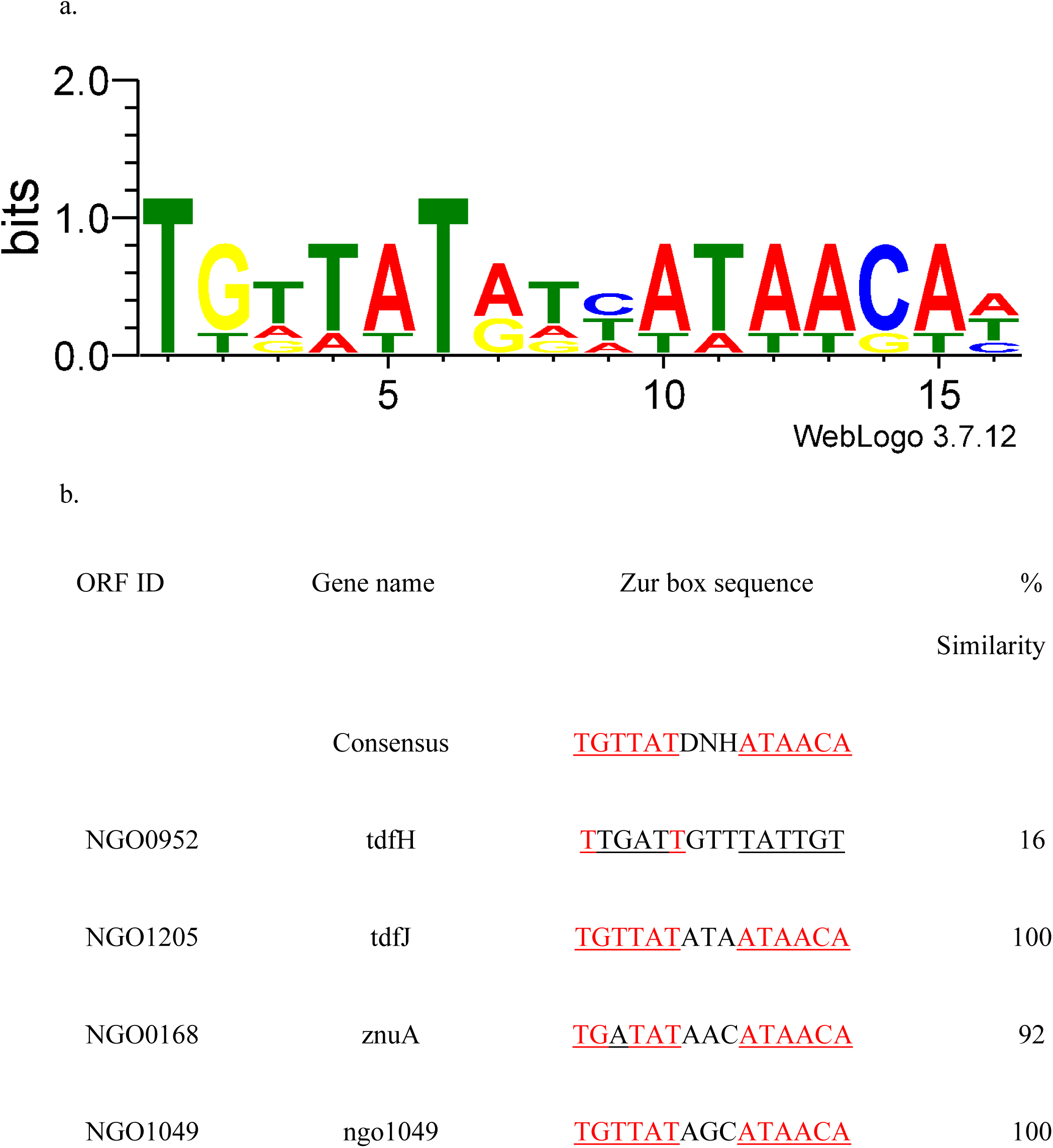
*tdfJ* and *ngo1049* harbor an exact match to the predicted Zur-box consensus sequence in their promoters. a) Graphical representation of the *Ngo* Zur-box was generated from predicted Zur binding locations on zinc-regulated promoters using WebLogo 3. b) A nucleotide sequence alignment of the predicted Zur-boxes in the promoters of the zinc-regulated genes in *Ngo*. The conserved palindromic repeats separated by three nucleotides in the middle are bolded and shown in red to signify complementarity to the Zur box consensus sequence.

**Figure 3.**
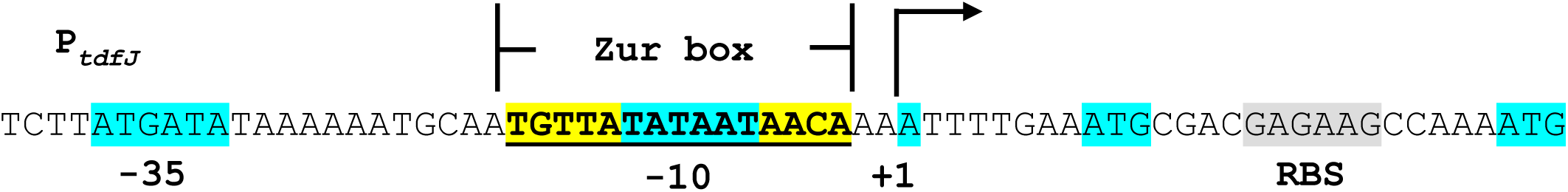
The *tdfJ* promoter elements based on 5’RACE analysis. The *tdfJ* promoter map shows the identified +1, predicted -10 and -35 regions and a Zur-box overlapping the -10 region. The two predicted initiator codon ATG, -10 and -35 and the +1 regions are highlighted in blue, and the Zur box palindrome is underlined and bolded in yellow. The canonical ribosome binding site (RBS) sequence is highlighted in grey.

### Two Fur-binding Sites on the tdfJ Promoter Result in Iron-dependent Induction of *tdfJ*

Along with the repression of TdfJ by zinc, *tdfJ* expression is induced in the presence of iron (10, 26). Yu. C, et al., showed that *tdfJ* expression was induced in wildtype cells in the presence of iron, but not in the fur mutant (26). We showed that this iron induction of TdfJ is further enhanced when zinc is absent (27). Additionally, our in-silico promoter analysis of *tdfJ* revealed two putative Fur-binding sites upstream of the -10 and -35 (Figure 4a). This site prediction was based on a compilation of known *Neisseria* Fur-box consensus sequences and our RNA sequencing data for iron regulation in FA1090 (26, 28, 29). This Fur-box consensus region was identified as a hexameric repeat of GATAAT-ATAATAATTATC-TTT (Figure 4b). On the *tdfJ* promoter (P_tdfJ_), the first predicted Fur-box site (Fur Box-1) lies about 58 base pairs upstream of the -35 region (Figure 4a). This region was included in the approximately 300 base pairs potential Fur-box location established by Yu et al., however a Fur binding affinity to P_tdfJ_ was not established from an EMSA (26). The second putative Fur-box site (Fur Box-2) was located overlapping the - 35, -10 regions and the Zur-box (Figure 4a). We hypothesized that the iron-induction of *tdfJ* is mediated through Fur and that Fur occupies both site-1 and 2 of P_tdfJ_, to activate gene expression when the promoter is de-repressed. To test this, we used the DNA-Protein Interaction ELISA approach (30). Briefly, biotinylated P_tdfJ_ was bound to streptavidin coated ELISA plates along with positive control P_tbpB_ and negative control P_rmpM_ DNA. Manganese loaded, purified recombinant His-tagged NgFur was added to the ELISA plate, for detection of DNA bound Fur using an anti-His antibody. The absorbance at 450nm (A_450_) was measured and binding affinity K_D_ was calculated with respect to the µg of Fur bound to the bound DNA. 10x excess unlabeled P_tdfJ_ competed with bound biotinylated P_tdfJ_ on the ELISA plate, resulting in reduced signal. Specific binding of Fur was calculated as: Specific Binding = µg of Fur bound to BIOTIN-tagged P_tdfJ_ - µg of Fur bound to 10x excess unlabeled P_tdfJ_. Fur was calculated to have a K_D_ of 2 for the positive control promoter, P_tbpB_, and a K_D_ of 6 for P_tdfJ_. A K_D_ was not able to be established for the negative control P_rmpM_ as the binding curve never achieved saturation at the concentrations tested (Figure 4c).

**Figure 4.**
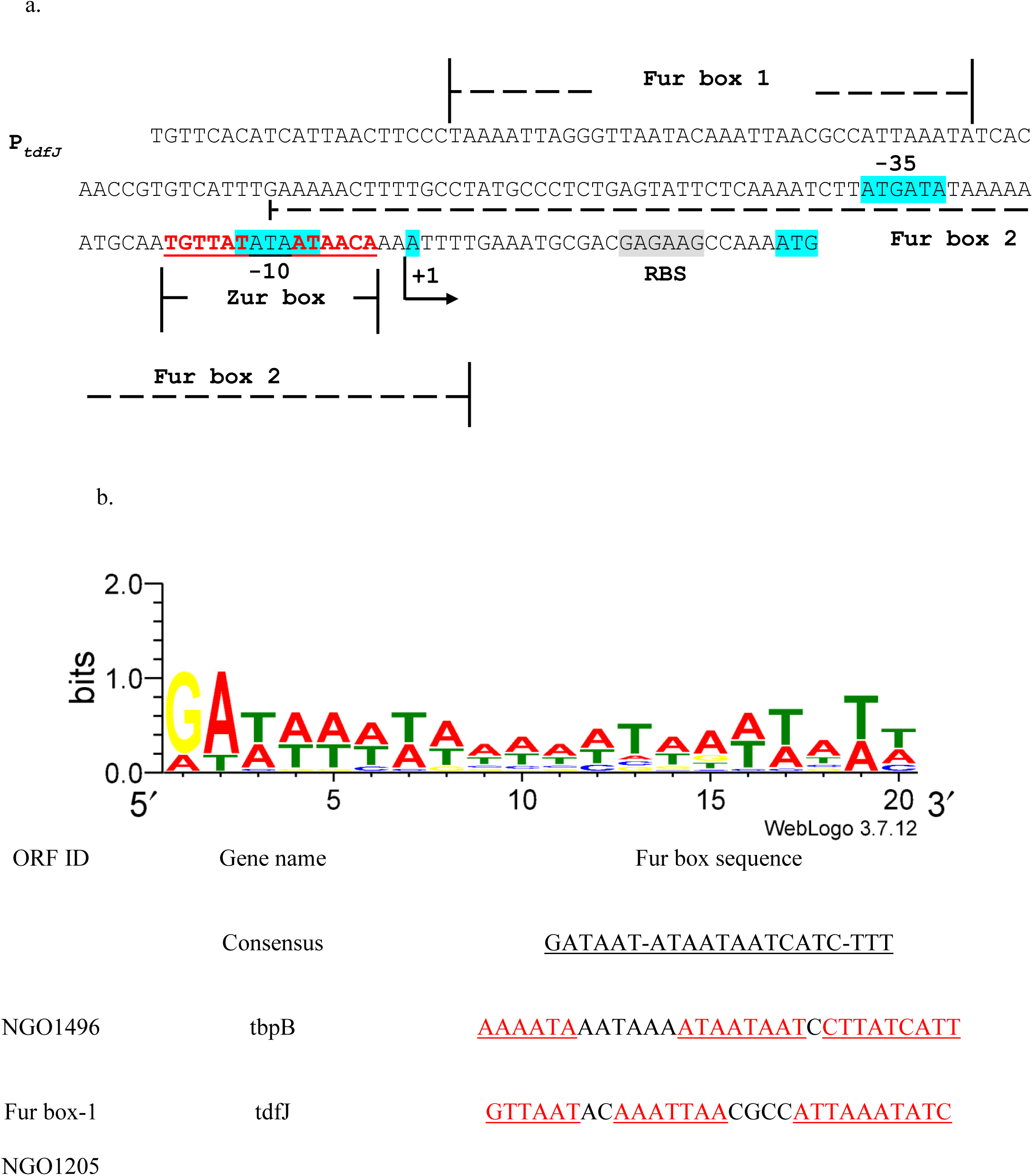
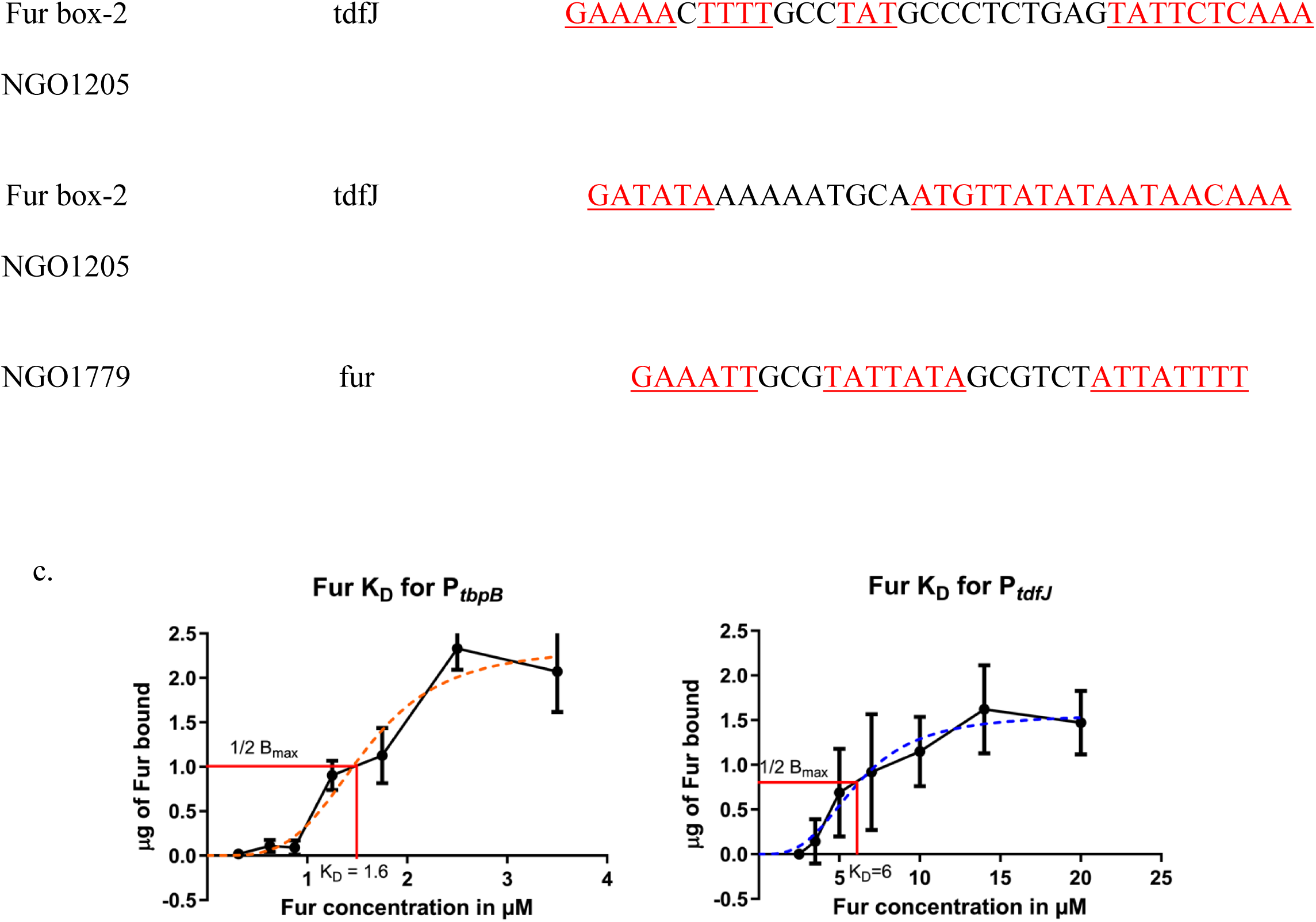
The *tdfJ* promoter contains two upstream Fur binding regions for iron regulation. a) The two locations of predicted Fur boxes upstream of *tdfJ* are labelled along with the identified tdfJ promoter elements. b) Graphical representation of the Fur-box of *Ngo* from predicted Fur binding locations was generated using WebLogo 3. The *Ngo* Fur box consensus sequence was determined from RNA Sequencing analysis of iron regulated promoters and consists of hexameric repeat regions. An alignment of the Fur box of Fur regulated promoters *tbpB* and fur are depicted along with the *tdfJ* promoter predicted Fur box. c) Fur binding affinity (K_D_) for positive control promoter P_tbpB_ and P_tdfJ_ promoter were calculated based on a linear regression model on Graphpad PRISM 5.0. The respective K_D_s are depicted as the slope intercept on the x-axis. The K_D_s are based on an average from three biological replicate ELISAs for Fur binding to P_tbpB_ and P_tdfJ_. P_rmpM_ was used as a negative control, not depicted here.

### Fur Binds the Specific Fur-box Consensus Sequence Regions 1 and 2 on P_tdfJ_

To establish the specificity of Fur for the two Fur-box sites on P_tdfJ_, we scrambled the predicted Fur-box sequences by changing the ATAAT-ATAATAATTAT-TTT repeats to a string of G’s and Cs (Figure 5a). We sequentially scrambled different portions of the 2 Fur-boxes and made 6 sets of scrambled P_tdfJ_, shown in Figure 5a from most to least scrambled. Biotinylated scrambled promoter sequences were bound to streptavidin ELISA plates along with biotinylated wildtype P_tdfJ_ and the level of 10 µM Fur binding measured. An increase in P_tdfJ_ scrambling correlated with a decrease in Fur binding absorbance values (A_450_), with the most scrambled P_tdfJ_ (both Fur-box 1 and 2) showing a significant loss in Fur binding (Figure 5b).

**Figure 5.**
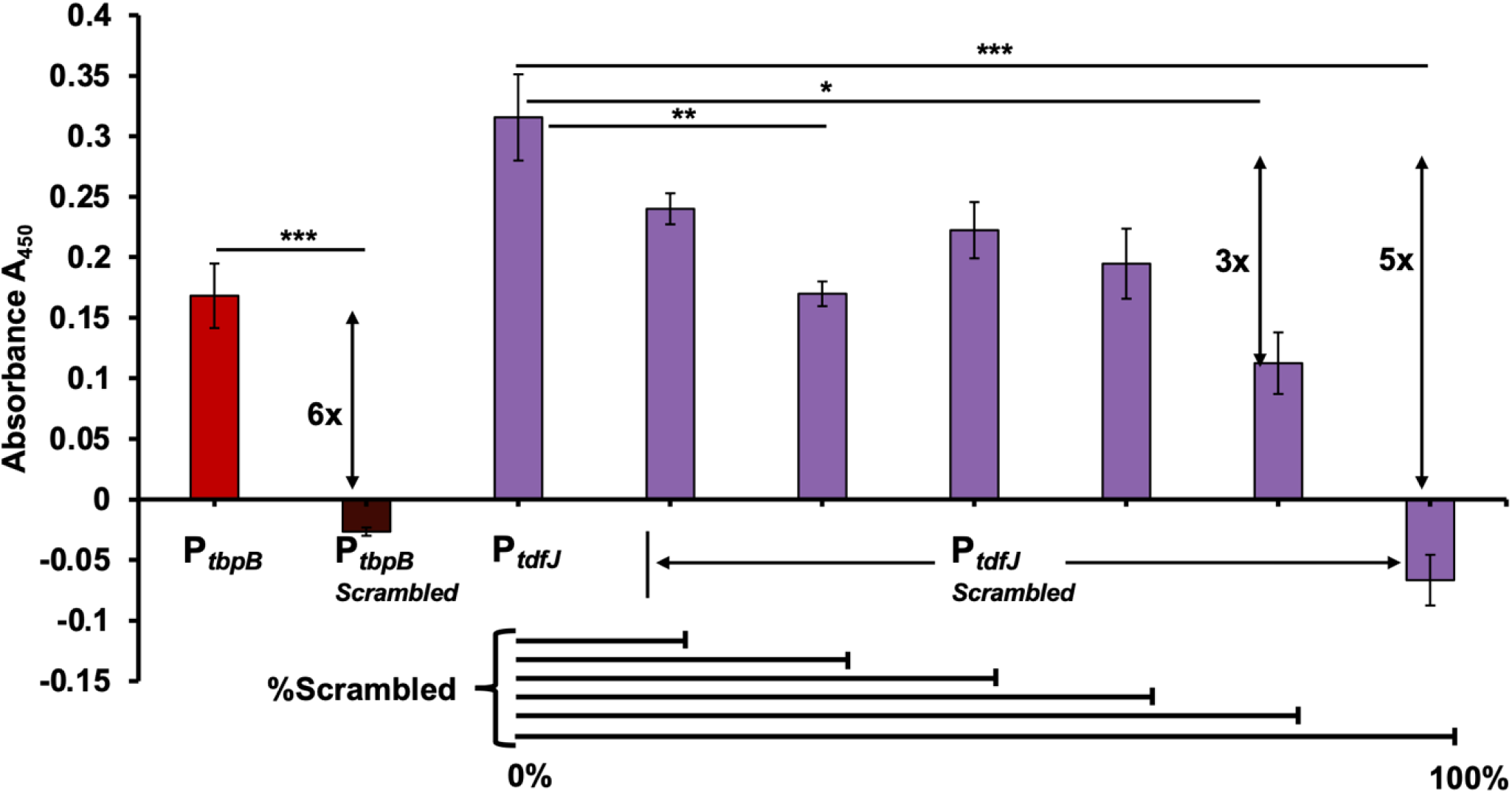
Fur binds specifically to the Fur-box hexameric repeat regions on P_tdfJ_. Fur-binding ELISA of wildtype and scrambled P_tbpB_ and P_tdfJ_ promoters. The A_450_ of Fur binding to promoters is on the x-axis, while promoter sequences are across the y-axis. Wild-type P_tbpB_ (light red) and scrambled P_tbpB_ (dark red) were used as positive and negative controls, respectively. Wild-type P_tdfJ_ promoter is in light green and scrambled P_tdfJ_ promoter sequences are denoted in dark green. Fold difference with respect to wildtype promoters is denoted by arrows. A student’s t test was performed for statistical analysis from three biological replicates. *, p<0.05; **, p<0.005; ***, p<0.0005.

## DISCUSSION

In this study, we show that *tdfJ*, *tdfH* and *znuA* are zinc-regulated by Zur at the transcriptional level, with *tdfJ* and *znuA* showing high fold-changes for gene expression. The gene expression results complement protein production. The gene expression levels in response to TPEN are similar in the wildtype and *zur* mutant, however for *tdfJ*, *znuA* and *ngo1049* the expression in the *zur* mutant in response to excess zinc appeared to be further de-repressed than from addition of TPEN alone. This was although not complementary in protein expression except for in Ngo1049. This suggests a tight control by Zur at the transcriptional level which is not fully derepressed upon addition of TPEN in this study. In theory we would have expected the zinc excess levels in the *zur* mutant to look like the TPEN treated condition. We should consider the effects of TPEN, as it not only restricts zinc but is able to chelate iron, and can disrupt cell membrane integrity, which could add to the lack of full de-repression by TPEN.

This study is the first to indicate the correct translational start for TdfJ, based on in silico and promoter characterization experiments, ending the long-standing debate on which ATG is utilized by *Ngo*. *tdfJ* promoter analysis established the presence of both Fur and Zur boxes helping explain a mechanism for dual regulation by iron and zinc. Through a novel DNA binding ELISA approach, we were able to show that Fur binds the *tdfJ* promoter with a specific affinity. We also show that Fur recognizes and binds more flexibly than possibly Zur. While Zur recognizes a much smaller Zur-box region, the Fur box is a much larger sequence of at least 30 bp and is spread across the *tdfJ* promoter (Figure 5a). TdfJ is unique among the TDTs for its ability to be dually regulated by both zinc and iron metal ions. The *Nme* homologue of *tdfJ*, *znuD* has also been shown to be induced by iron and repressed by zinc, with a speculation of being Fur- and Zur-mediated respectively (31). Analysis of the *znuD* putative promoter elements implicates a striking match to the *tdfJ* promoter characterized in this study. The mechanism that Fur utilizes to activate gene expression here has not been characterized, however, based on existing evidence Fur can activate *tdfJ* expression through a direct mechanism involving Fur binding to P_tdfJ_ (26). The direct mechanism may involve competition between Fur and a repressor, for the same site on the promoter. This has been previously demonstrated in the activation of *Ngo norB*, by Fur, where Fur competes with a repressor, ArsR for binding the Fur-box, thereby resulting in de-repression and increase in expression of *norB* (32). However, this mechanism in the context of P_tdfJ_ has not been considered because Fur, the activator, and Zur, the repressor, are not competing for the same site on the promoter (Figure 5a). Based on the location of the Fur-box, and evidence from other studies, the Fur-box is at a position optimal for RNA polymerase recruitment (26, 33, 34). Zur as a repressor is not bound when zinc is absent, which is a condition necessary for *tdfJ* expression (13, 16). TdfJ is unique among the TDTs for its ability to be dually regulated by both zinc and iron metal ions. The *Nme* homologue of *tdfJ*, *znuD* has also been shown to be induced by iron and repressed by zinc, with a speculation of being Fur- and Zur-mediated respectively (31). Analysis of the *znuD* putative promoter elements implicates a striking match to the *tdfJ* promoter characterized in this study. The mechanism that Fur utilizes to activate gene expression here has not been characterized, however, based on existing evidence Fur can activate *tdfJ* expression through a direct mechanism involving Fur binding to P_tdfJ_ (26). The direct mechanism may involve competition between Fur and a repressor, for the same site on the promoter. This has been previously demonstrated in the activation of *Ngo norB*, by Fur, where Fur competes with a repressor, ArsR for binding the Fur-box, thereby resulting in de-repression and increase in expression of *norB* (32). However, this mechanism in the context of P_tdfJ_ has not been considered because Fur, the activator, and Zur, the repressor, are not competing for the same site on the promoter (Figure 5a). Based on the location of the Fur-box, and evidence from other studies, the Fur-box is at a position optimal for RNA polymerase recruitment (26, 33, 34). From the context of both Fur and Zur regulating *tdfJ*, we need to understand if they would act at the same time and which regulator has a greater affinity for the promoter. This would depend on the signal, which is the presence or absence of their cognate metal ions Fe^2+^ and Zn^2+^ respectively. Since Fur recognizes a larger area spanning from the upstream of the -35 region to the -10 region (denoted Fur box 1 and 2) (Figure 5a), sterically it would not be possible for Zur also to be bound at the same moment. We hypothesize that when Zur is bound in the presence of Zn^2+^, the interaction between Zur and the *tdfJ* Zur box is of greater affinity, preventing Fur-Fe^2+^ from displacing Zur to act on *tdfJ*. However, the absence of Zn^2+^ and therefore Zur would allow room for Fur to bind in the presence of Fe^2+^ to induce *tdfJ* expression by recruiting RNA Polymerase. This hypothesis needs to be tested experimentally to determine: 1) the specificity and affinity of Zur to the *tdfJ* promoter and 2) competition between Fur and Zur for the *tdfJ* promoter which acting at the same time at various Mn^2+^ and Zn^2+^ concentrations. Preliminary efforts have been taken to address the binding affinity and specificity of Zur to P_tdfJ_, however, several obstacles were faced with obtaining a stable form of purified recombinant NgZur for binding assays. Protein sequence analysis for *Ngo* Zur showed strain specific difference in the length of Zur between FA1090 and FA19 with the latter from attaining stability during crystallization efforts. Protein crystallization efforts are currently being carried out at Purdue University’s Department of Biological Sciences by Dr. Nicholas Noinaj and Swati Mundre, to obtain a recombinant form of FA1090 NgZur that is stable when purified to use in our DNA binding ELISAs. Zur as a repressor is not bound when zinc is absent, which is a condition necessary for *tdfJ* expression (13, 16).

In a recent study characterizing the Zur regulon in Neisseria meningitidis (*Nme*), *Nme* Zur differentially regulated genes of the Zur regulon, by binding to a Zur-box that was identical to the *Ngo* Zur-box identified here (Figure 2) (16). *Nme* homologues of *tdfJ* (*nmb0964*) and *ngo1049* (*nmb1475*) also showed perfect matches to their predicted *Nme* Zur-box and showed high fold-changes for gene expression in the presence of TPEN. This conservation of Zur-boxes among the pathogenic *Neisseria* species, is interesting and illustrates the importance of evolutionary conservation to maintain zinc homeostasis. However, whether this is specific to pathogenic *Neisseria* has not been probed into. Since several of the host adaptations have been conserved across both commensals and pathogenic species, it would not be surprising if Zur boxes were also conserved among the *Neisseria* commensals. This conservation of Zur-boxes has been investigated across several β-proteobacteria, including *Bacillus subtillis*, *Yersinia pestis*, *Pseudomonas aeruginosa*, *Escherichia coli*, *Acinetobacter baumanii* and *Salmonella typhimurium* and the conserved sequence was identified to be <u>GAAATGTTATA</u>-N-<u>TATAACATTTC</u> (15). All these studies have showed the potential of Zur in pathogensis due to its importance in maintanence of zinc homeostasis of several virulence genes, in *Ngo* these important zinc homeostasis virulence genes are *tdfH*, *tdfJ* and *znuA*.

Zur maintains zinc homeostasis by tightly controlling the level of expression of the genes it regulates. Zinc-bound Zur dimerizes and represses transcription of genes by binding to a consensus Zur-box on the promoter of the gene, thereby blocking RNA polymerase transcription initiation (17). In this study, we demonstrate a possible correlation between a consensus Zur-box and higher a higher fold-change for gene expression. *tdfJ*, *znuA* and *ngo1049* with a 90-100% match to the Zur box also show a 24, 38 and 46-fold-change for expression respectively in the absence of zinc, however *tdfH* with a 16% match to the Zur-box consensus shows only a fold-change of 3. (Figures 1 and 2). Therefore, the strength of gene regulation by *Ngo* Zur is dependent on the accuracy and length of the palindromic consensus sequence. Further, TdfJ has been identified as an ideal vaccine antigen for the prevention of gonorrhea (10, 35). Its, tight transcriptional regulation by Zur in response to zinc, demonstrate its potential to be a molecular tool that will allow further investigation of its role during infection and enhance the understanding behind creating an efficacious vaccine. This requires further characterization of the *tdfJ* promoter elements that allow it to be regulated by Zur and zinc.

It is also important to consider the physiological relevance of the existence of an environment that will allow for TdfJ expression particularly, that is iron enriched, and zinc deprived. To answer this question, we would need to scope the availability of zinc and iron, or their lack thereof, for the expression of *tdfJ*. Zinc is primarily secreted by the prostate and is abundant in the male reproductive organs and human semen as compared to other body tissues and secretions (36). In contrast, much less is known about the bioavailability of zinc in the female reproductive tract, but is known to be more zinc starved with vaginal fluid comprising of no zinc in daily secretions (37). S100A7 and calprotectin zinc-sequestering proteins, are also enriched in the ectocervix of the female reproductive tract, causing low levels of zinc (38). Overall, the female reproductive tract is more zinc-starved compared to the male reproductive tract, making the female reproductive tract an ideal environment for the expression of *tdfJ* and other Zur-repressed genes. In addition to low levels of zinc, iron enrichment occurs in the female vaginal canal during the menstrual cycle when tissue and red blood cells lyse to release heme and hemoglobin that are sources of iron. This provides a conducive zinc-starved iron-rich surrounding for enhanced *tdfJ* expression. It is, however, unknown if TdfJ plays a role in the asymptomatic nature of infections predominately seen in women. Preliminary studies on the involvement of TDTs in invasion of ME180 cervical epithelial cells showed promising results for TdfJ as a potential invasion, although the study concluded that only TdfF was important during invasion. There is scope for further analysis of these results and can be revisited to explain TdfJ s role during infection of cervical versus urogenital epithelium (39). There is much to be understood on the nutrient metal bioavailability during gonococcal pathogenesis and the combined effect of multifactorial players. Using the *tdfJ* promoter as a molecular tool to access the environmental differences between the male and female genital tracts in terms of iron and zinc, will expand the scope and knowledge on these multifactorial players and help parse out the mechanisms of gene regulation that take place during gonococcal pathogenesis.

Collectively, our study is the first to lay out a dual mechanism of regulation of TdfJ, by two distinct metal ions and regulators Fur and Zur. Future studies testing the *tdfJ* promoter as a zinc and iron sensitive reporter in different pathological environments including neutrophils, where *Ngo* survive phagocytosis, epithelial cells, and transgenic mice infection models, will be valuable to the field and aid in deciphering targeted treatment options and outcomes.

## Materials and Methods

### Bacterial Strains and their growth conditions

Strains and plasmids used in this study are listed in Table 1. Plasmids were propagated in *Escherichia coli* strains TOP10, DH5alpha or BL21 DE3 pLysE. *E. coli* strains were cultured in Luria-Betrani (LB) broth media supplemented with antibiotics carbenicillin (100 µg/mL), kanamycin (50 µg/mL) or erythromycin (34 µg/mL). *Neisseria gonorrhoeae* (*Ngo*) WT and mutant strains were routinely maintained on GC medium (Difco) base agar plates with Kellogg’s supplement and 12 µM Fe(NO_3_)_3_ (GCB Plates) at 36°C and 5% CO_2_ atmospheric conditions .

**Table 1.**
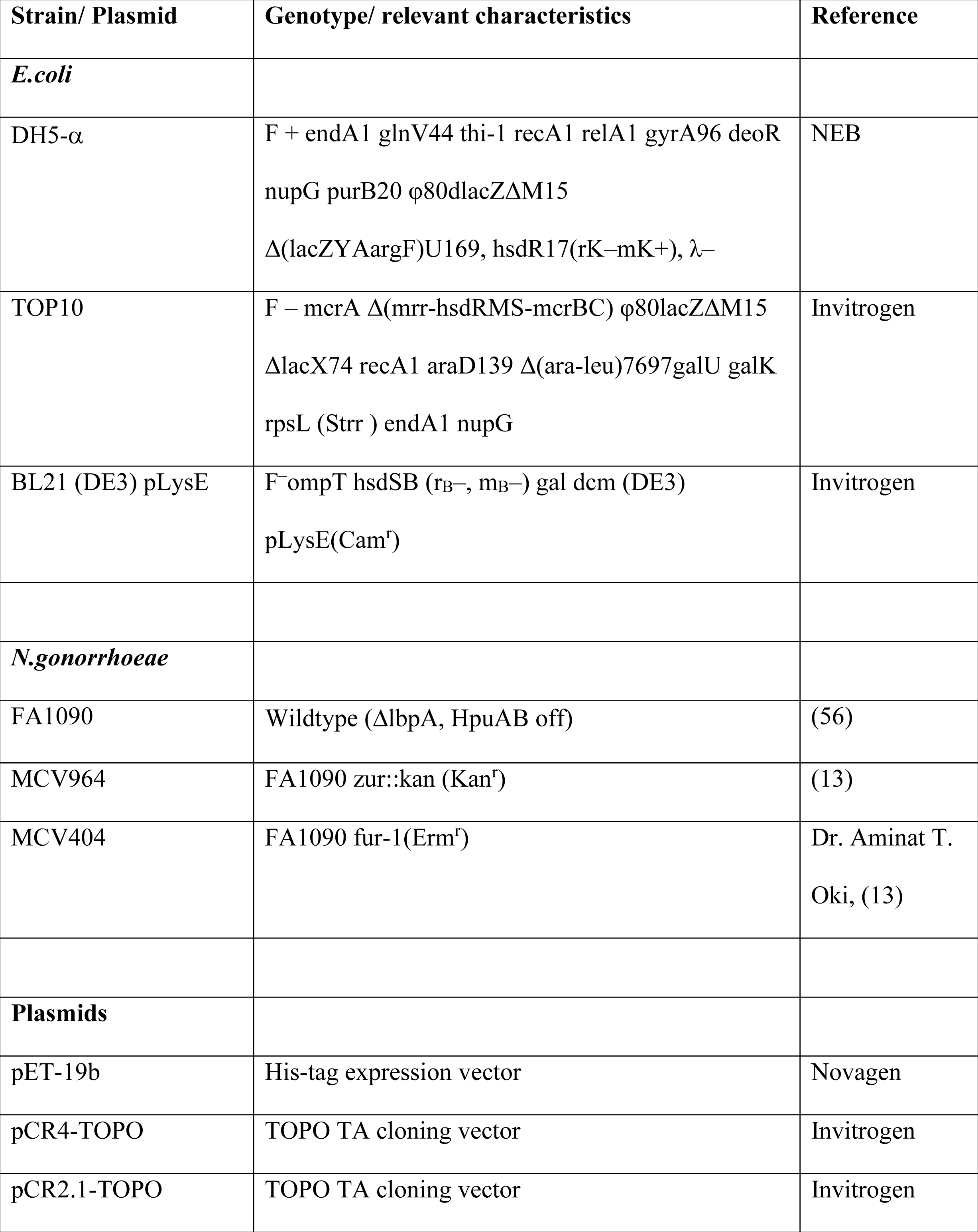
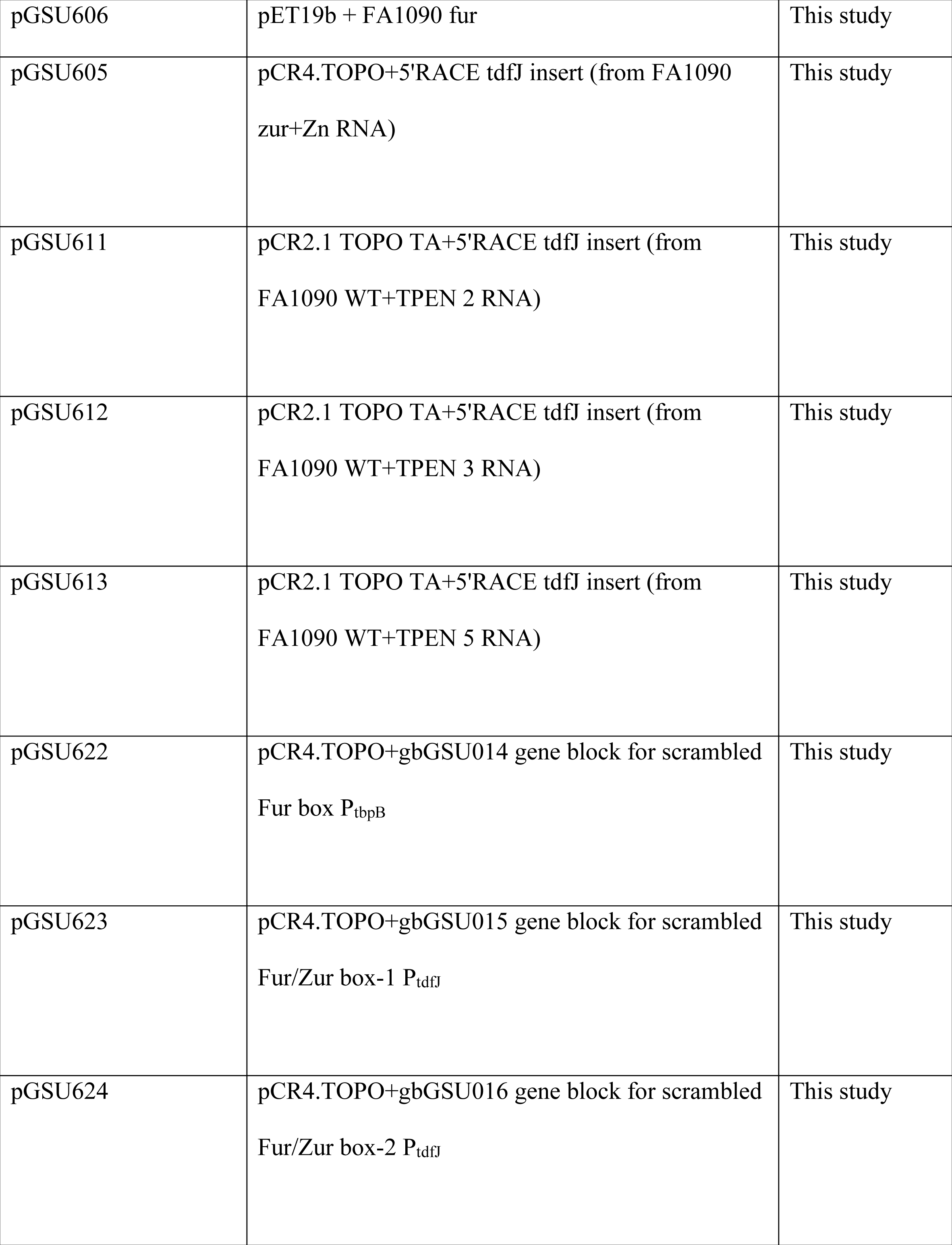
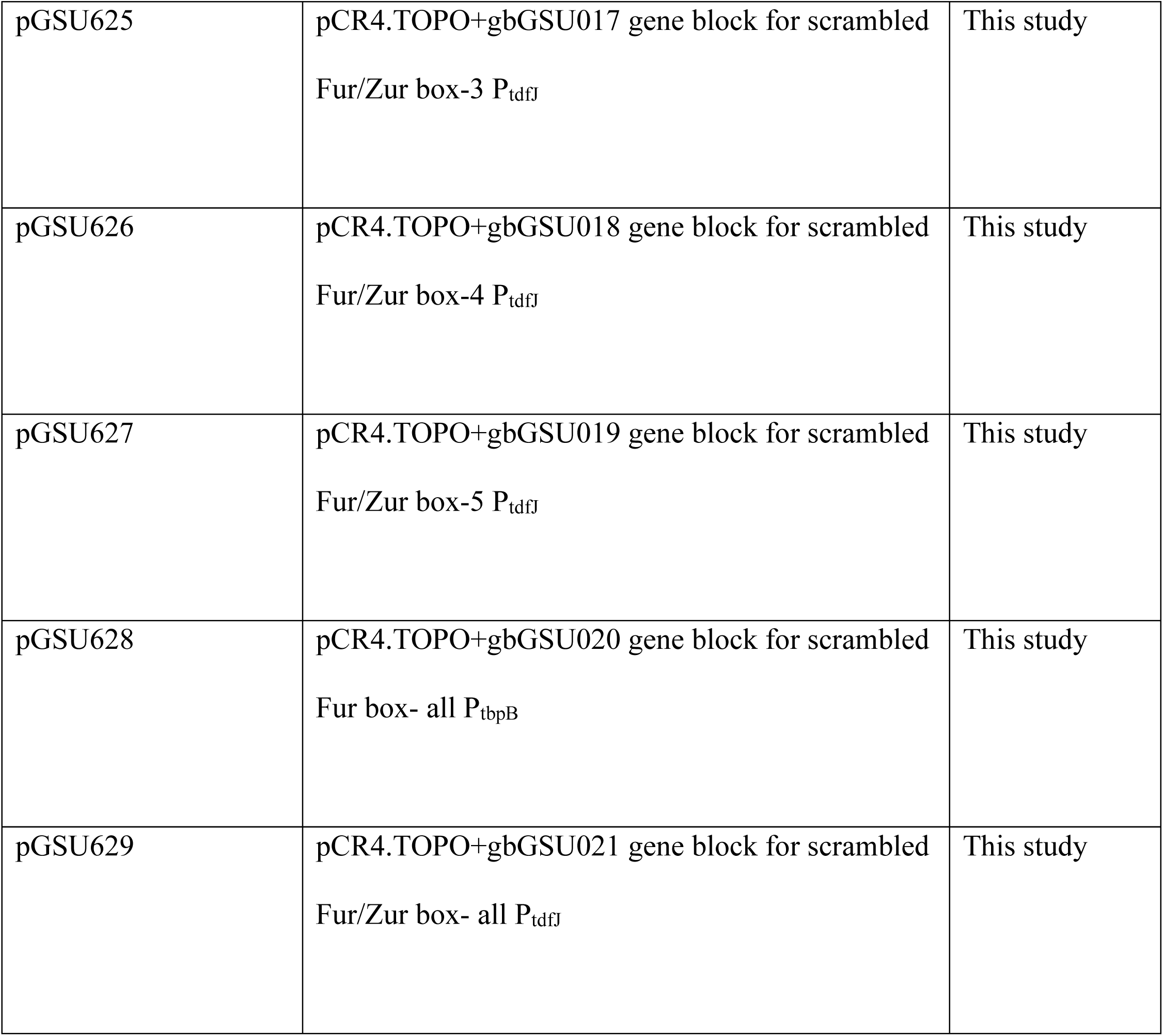
Strains and plasmids used in this study.

### *Ngo* Growth Conditions for Zinc and Zur Regulation

*Ngo* strains FA1090 WT and MCV964 (FA1090 *zur*::*kan*) (Table 1) (13), were grown under zinc-restricted conditions as previously described (40). Briefly, non-piliated colonies were selected and passaged onto GCB plates containing 5 µM N,N,N’ N’ -tetrakis-(2-pyridylmethyl)-ethylenediamine (TPEN) (Sigma), a zinc-specific, to deplete any internal zinc stores of Ngo. Single non-piliated gonococcal cells from GCB TPEN plates, were used inoculated into 50 mL chelex-treated chemically defined media (CDM) at an initial OD_600_ of ≅ 0.9. The strains were grown at 36°C and 5% CO_2_ with vigorous shaking at 225 RPM in acid-washed baffled flasks and allowed to grow exponentially. After doubling of OD_600_ (approx. 1.5 to 2 hours), the CDM was supplemented with either 20µM Zn(SO4)2 for excess zinc condition or 7 µM TPEN for zinc-deplete condition. Both strains were allowed to grow to mid or late log phase (about 2.5 to 3 hours), with the OD_600_ being measured every 30 min to 1 hour, to track the growth curve. Whole cell lysates were collected by normalizing the OD to 100,00KU (Klett units) at 600nm. The remaining sample was used for RNA isolation, for subsequent RT-qPCR or 5’RACE.

### RNA Isolation and Processing for Gene Regulation and Promoter Analysis

RNA was harvested from the 50 mL zinc-restricted growth cultures, at mid-to late-log phase. The cultures were pelleted and resuspended in Nucleoprotect RNA (Macherey-Nagel) according to manufacturer’s protocol (41). After complete permeation, samples were centrifuged at 10,000 rpm for 2 minutes to remove RNA protect and pellets were stored at -80C until RNA extraction. The stored pellets were thawed on ice on the day of RNA isolation under a laminar flow hood. The hood was cleaned with 10% bleach, 70% ethanol, DNA ZAP (Invitrogen Cat: AM9890) reagent and water prior to use. The unit was sterilized under UV for at least 30 mins before RNA isolation. RNA isolation was performed using the NucleoMag RNA isolation Kit (Macherey Nagel) in a semi-automated system using the Eppendorf EpMotion 5073 liquid handler, according to manufacturer’s instructions (42). User intervention was added at the clear lysate step to allow for external centrifugation of the samples to generate a lysate. The bench spaces were cleaned thoroughly with DNA Zap, RNase Zap (Invitrogen Cat:AM9780) and 5% bleach. Samples pellets were thawed on ice and placed in the EpMotion benchtop along with the RNA isolation kit reagents, DNA and RNAse-free tips, and reservoirs, at their respective designated positions, programmed in the EpMotion software. After isolation, RNA samples were further treated with TURBO DNase (Invitrogen Cat: AM2238) according to manufacturer’s protocol (43). cDNA was generated using the Superscript IV First-Strand Synthesis System (Invitrogen Cat: 18091050), and quality of samples were validated through + and – reverse transcriptase controls and bleach agarose gel electrophoresis (44).

### Quality Control for RNA and cDNA

RNA isolated by EpMotion was treated with TURBO DNase (as mentioned in the previous step) after a first round of concentration measurement. Approximately 2 µg of total RNA (200 ng/µL) isolated from each condition and each replicate was treated with TURBO DNase according to manufacturer’s protocol (43). A small portion of the DNase-treated RNA diluted to a final concentration of 25 ng/µL was used to examine the quality of the RNA via a PCR. PCR reactions with 5 ng RNA per reaction was setup using the GoTaq 2x mastermix and RNAse-DNA-free water for *rmpM* expression. The PCR reaction was run for 35 cycles in a BioRad thermocycler and the products were analyzed using a 2% agarose gel electrophoresis in 1XTris-acetate EDTA (TAE) buffer. The gel was run at constant voltage of 135V and imaged on a Chemidoc under UV trans illumination. Presence of bands on the gel from the PCR products indicated contamination with DNA resulting in further cleanup of RNA. Clear gel with no PCR products were desired results and positive control FA1090 genomic DNA and negative no template controls were used to make decisions on quality. Similarly, cDNA products of clean RNA + and – reverse transcriptase (RT) were subject to quality control PCR with rmpM primers oGUS195 and oGSU196. These products were analyzed on a Chemidoc after agarose gel electrophoresis and absence of products in – RT samples were expected for clean RNA.

DNase-treated RNA samples were further controlled for their quality using a bleach gel electrophoresis method for RNA Seq sample preparation (45). Total DNase-treated RNA at a concentration of 500 ng to 1 µg was diluted with 6x DNA loading dye and loaded onto a 5% bleach 1% agarose gel and run in 1xTAE at 135 V. The gel was imaged using a Chemidoc under a UV transilluminator to visualize 5S, 16S and 23SrRNAs at 120bp, 14kb and 30 Kb respectively.

### Primer Design and Efficiency Testing

Prior to RT-qPCR, primers (Table 2) were carefully designed to be less than 25 bp in length, amplify regions no greater than 200 bp and tested to have primer efficiencies between 90% and 95%. The primer sequence stability, hairpins and ΔG values were analyzed on the IDT website prior to placing primer orders for testing (46). Primer efficiencies were tested in gradient concentrations of primer ranging from 10 µM to 40 µM concentrations and gradient concentrations of genomic FA1090 DNA from 1 µg serially diluted by a factor of 10. PCR was set up for each primer set and each primer concentration with the SYBR Hi-ROX Sensimix (Bioline Cat: QT60505) along with melt curve data to check for primer specificity. PCR program was setup in the CFX96 BioRad thermocycler and standard curves for the range of DNA concentrations were analyzed for primer efficiency. Primers with annealing temperatures at 60°C were chosen for each gene, to match the annealing temperature of the rmpM reference gene primer set that was determined after primer efficiency tests. Primers and concentrations that yielded efficiencies between 90 and 95% were chosen for RT-qPCR analysis of target genes. Standard curve for these efficiency tests were plotted using the BioRad CFX96 software.

**Table 2.**
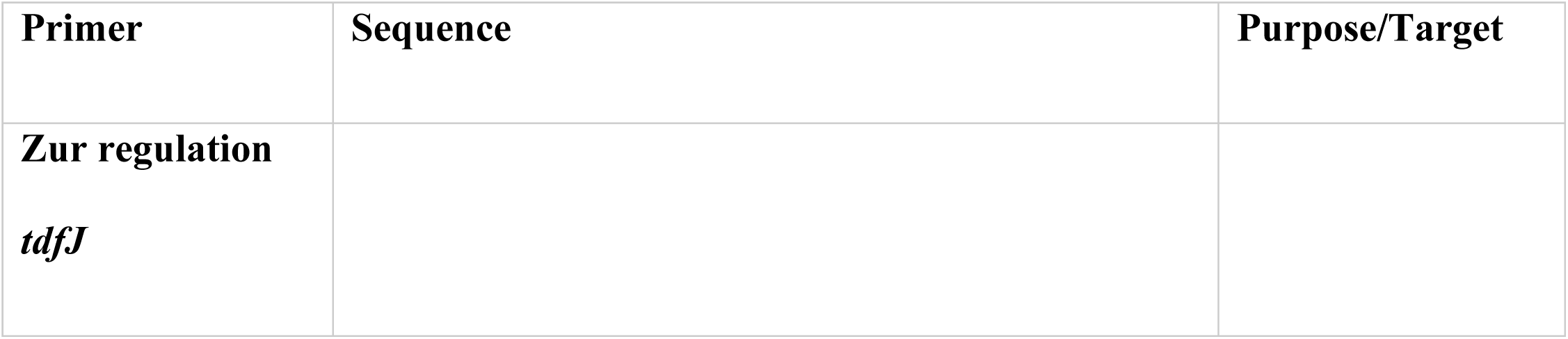
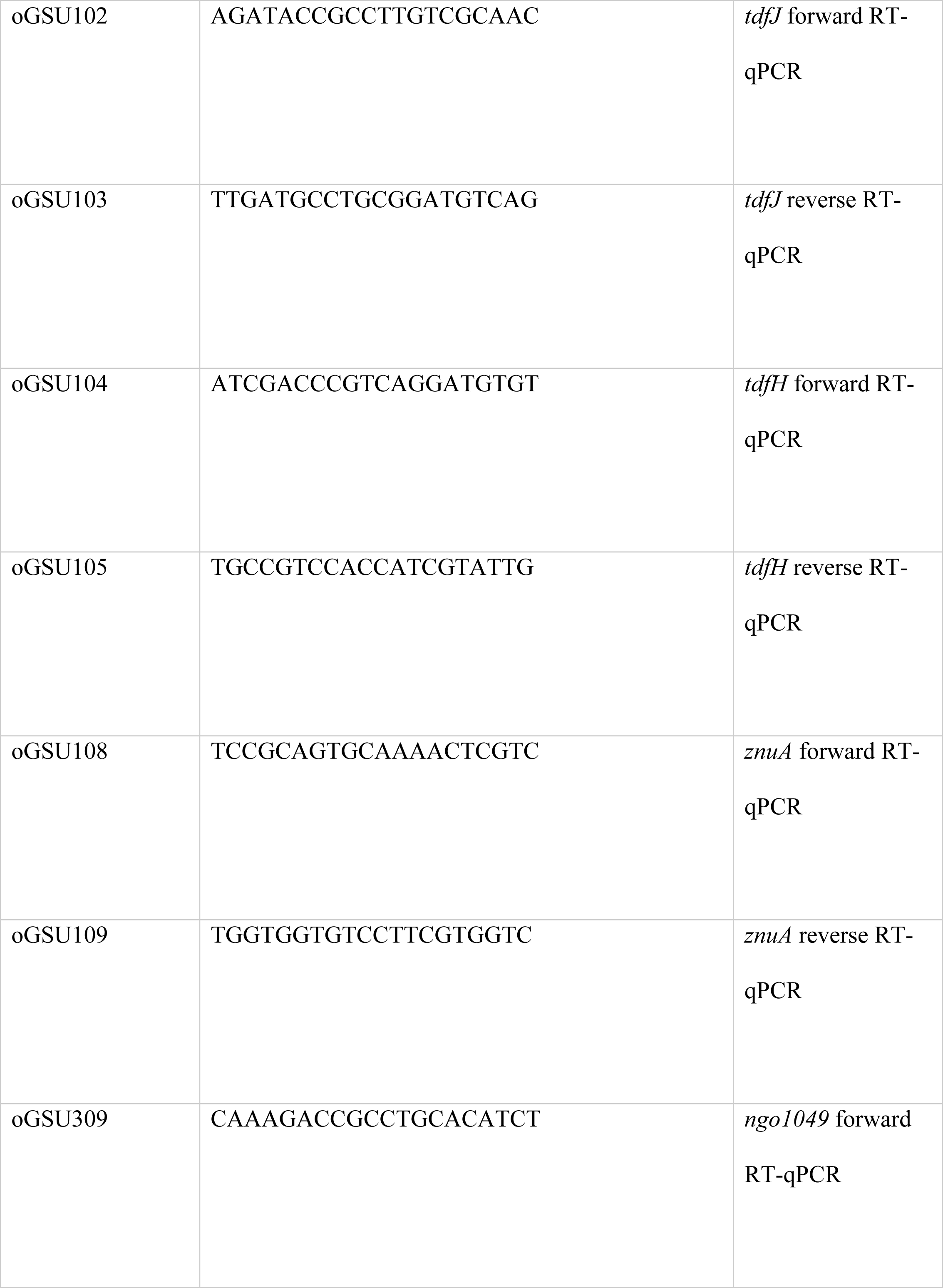
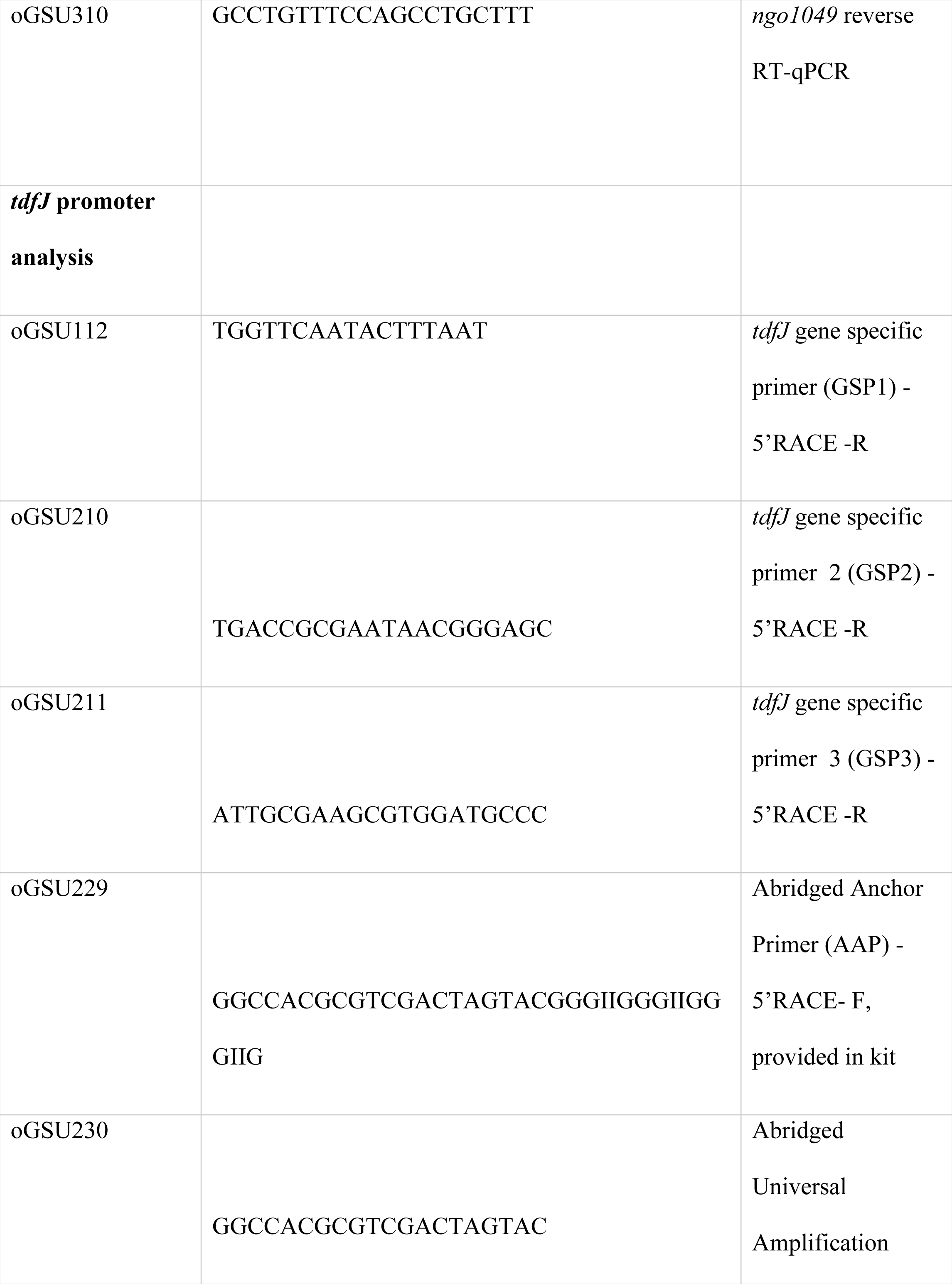
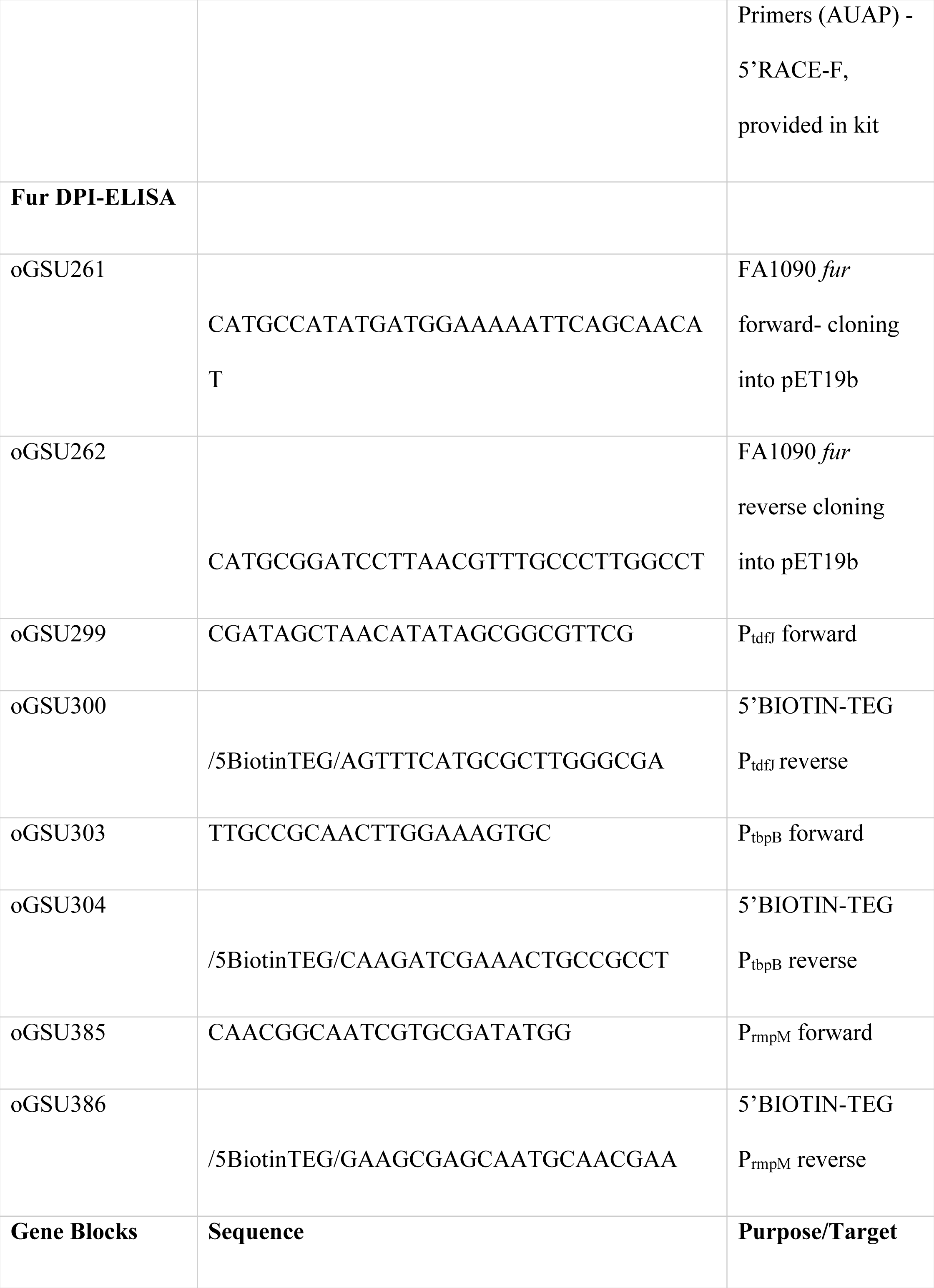
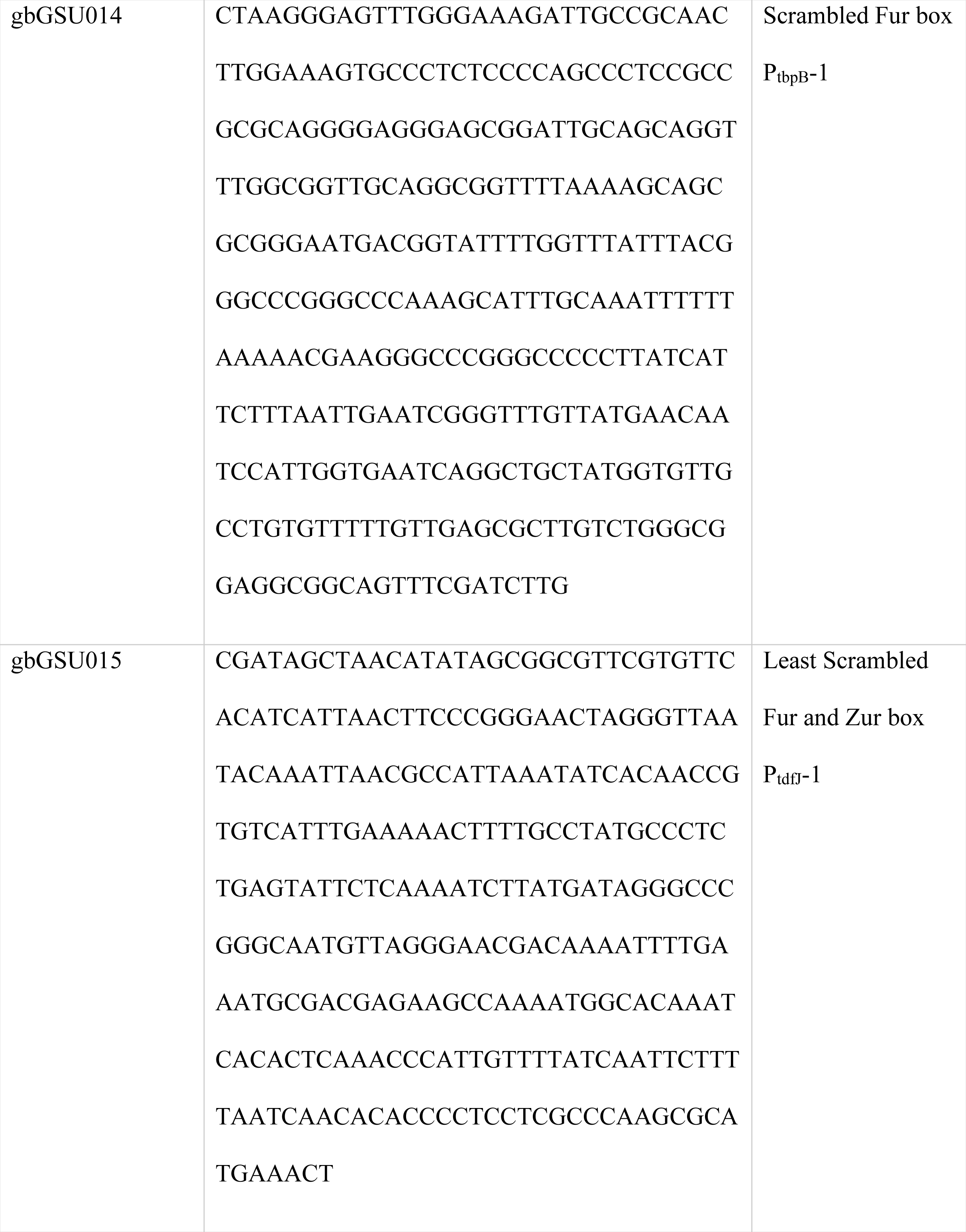
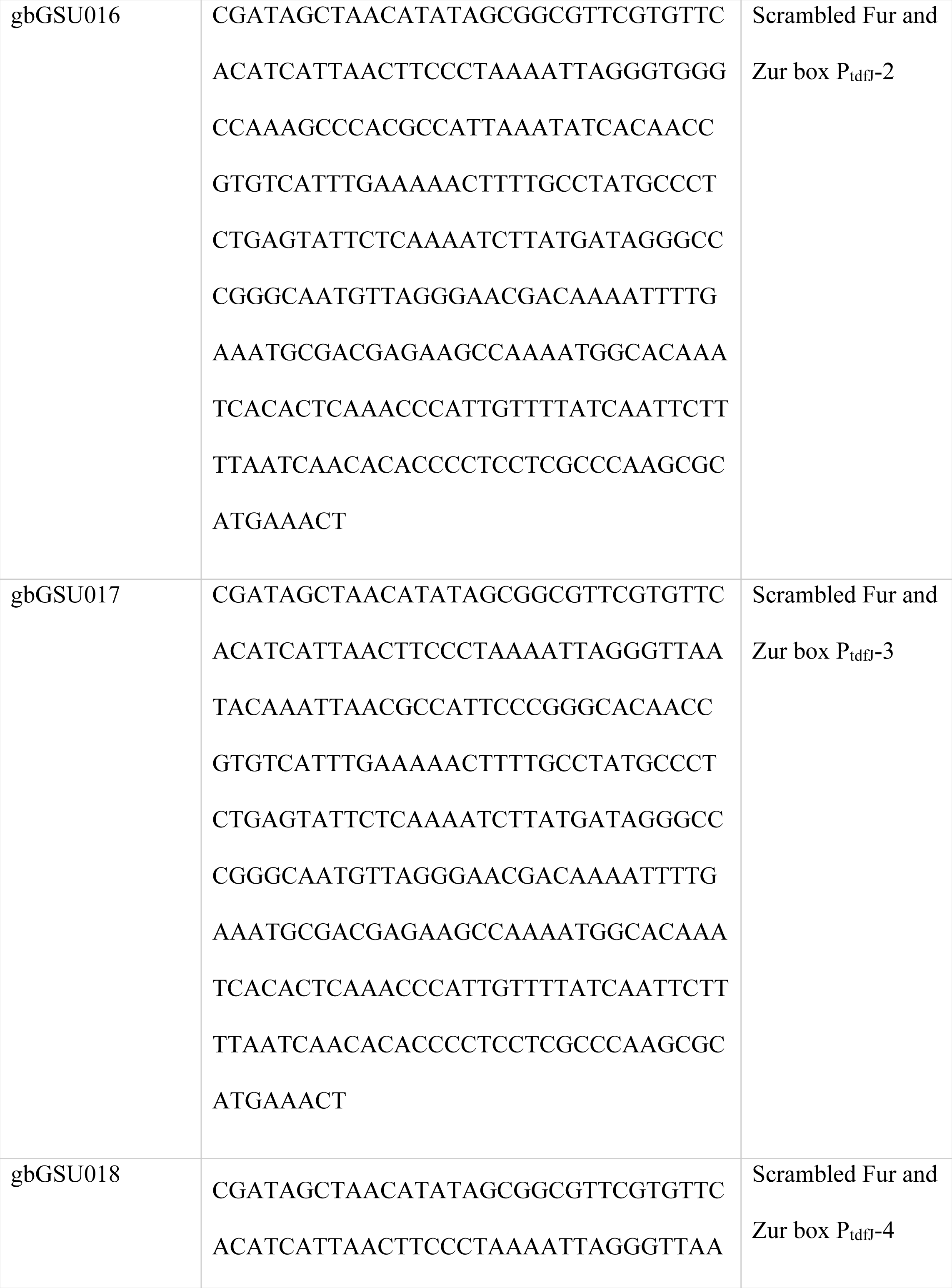
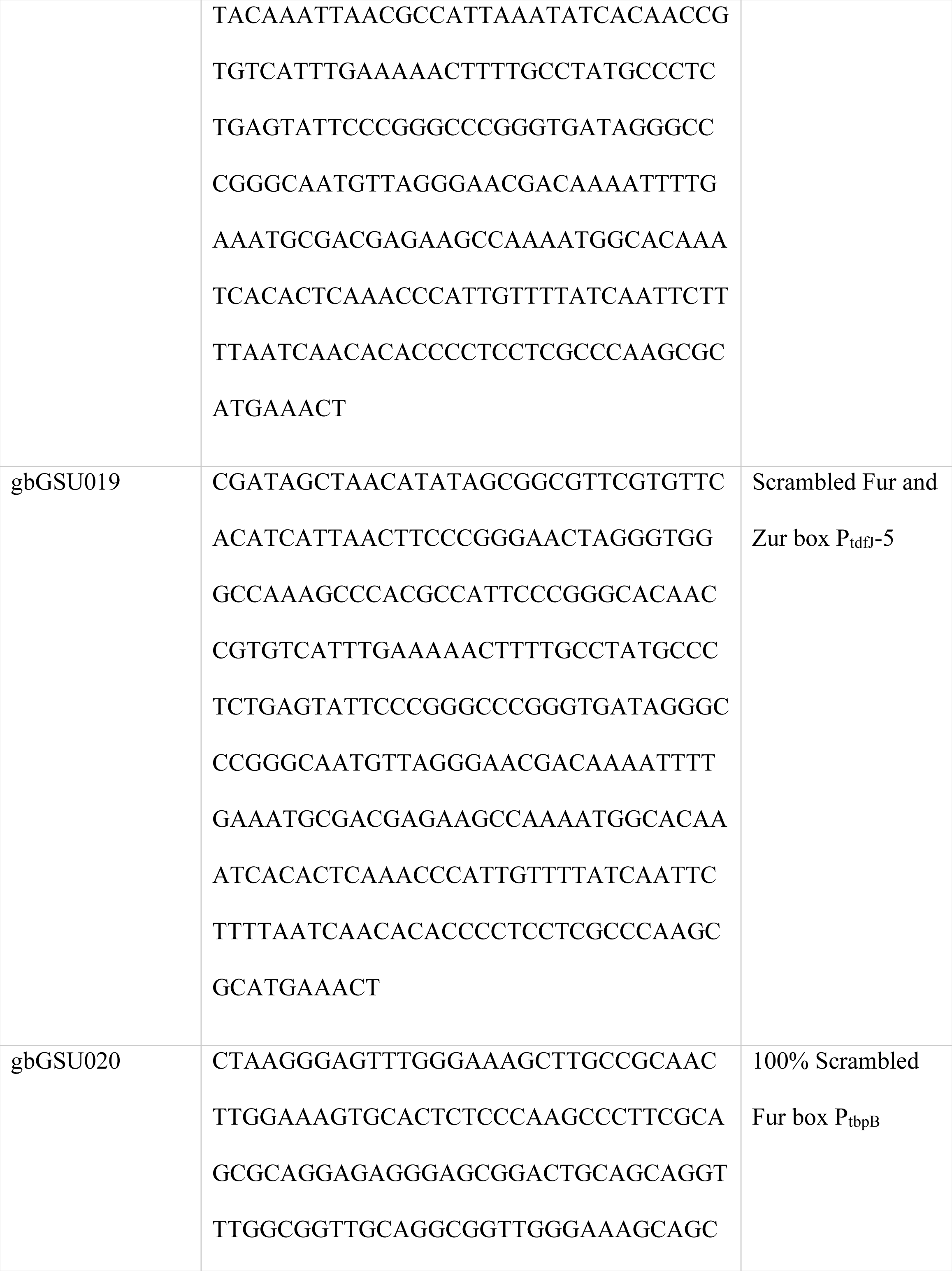
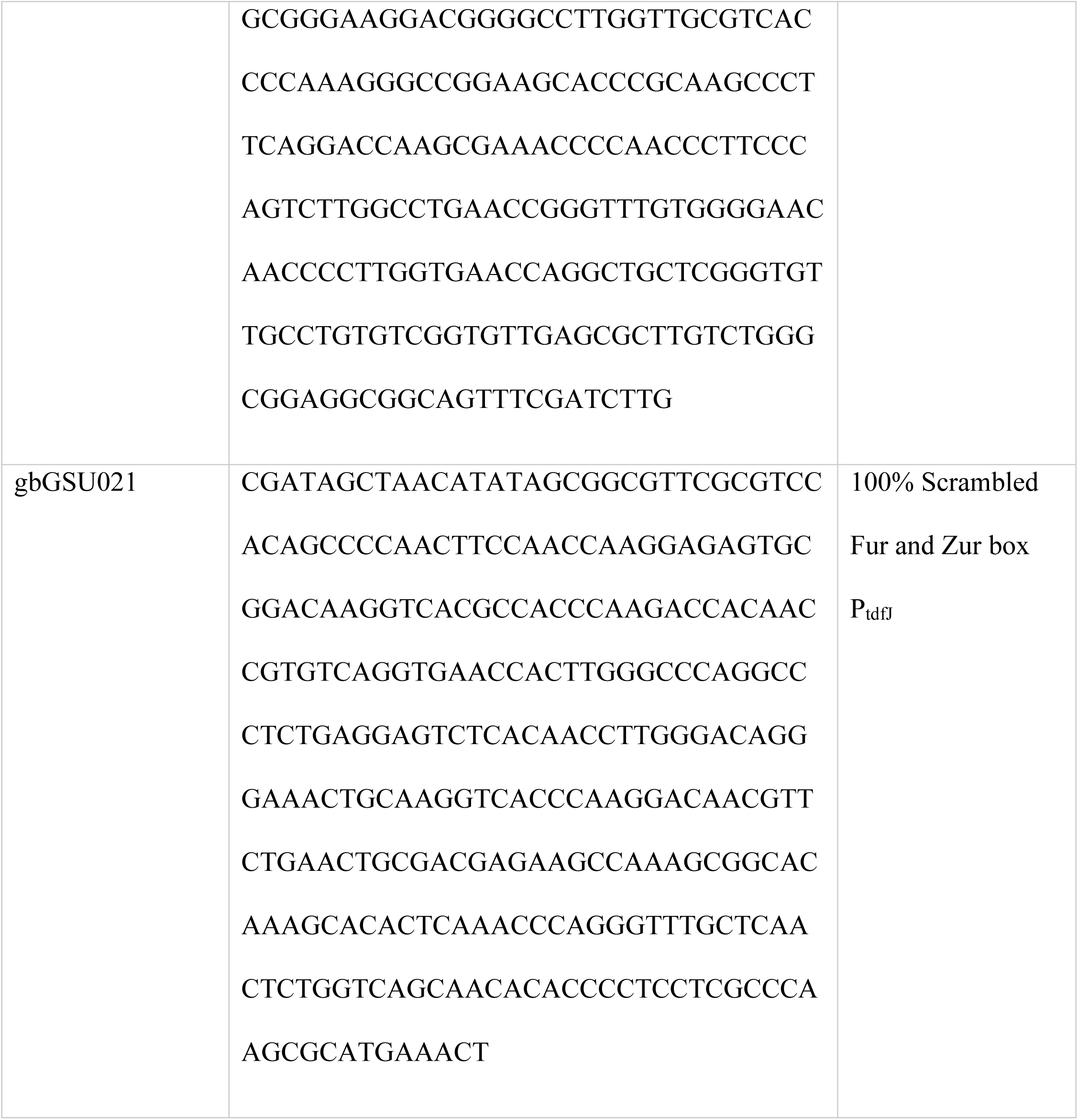
Primers and Gene Blocks used in this study.

### RT-qPCR for Measuring Target Gene Expression

RT-qPCR was performed with the SYBR Hi-ROX SensiMix (Bioline Cat: QT60505) polymerase on a Bio-Rad CFX96 thermocycler. RT-qPCR was set up for 35 cycles and normalized expression of each target gene was calculated in biological and technical triplicates. The specificity of the primers was checked by a melt curve (95°C for 1 min, 55°C for 1 min and an increment in temperature to 95°C at 0.5°C/s rate for 10 s). Gene expression was normalized to the expression of the housekeeping reference gene *rmpM*. Expression was calculated using ΔCq values with the E = 2^ΔCq^ formula where E is the expression and ΔCq is the normalized threshold cycle value for each gene (47).

### 5’ Rapid Amplification of cDNA Ends (5’ RACE) for Identification of the Transcriptional Start Site

The *tdfJ* transcript 5’ end was identified using the 5’ RACE Version 2.0 Kit according to manufacturer’s protocol (24). Two individually isolated RNA samples from either the FA1090 WT+ TPEN or FA1090 *zur*+ TPEN sample, were used to determine the transcriptional start site. *tdfJ* gene specific primer 1 oGSU112 was used for making gene specific cDNA (Table 2). Gene-specific primers oGSU210 and oGSU211 (Table 2) were used for the amplification of 5′ RACE gene-specific PCR products with AAP provided in the kit (Table 2). Gene specific PCR was performed on a Bio-Rad thermocycler using the Platinum Taq DNA polymerase (Invitrogen Cat: 15966005) and 10 mM dNTPs (48). The PCR products were cleaned and concentrated before cloning into the pCR2.1-TOPO TA vector (Invitrogen Cat: K457502) and sequenced at Genewiz (49). Sequence analysis was performed by aligning sequences from at least three different plasmids, after cloning the RACE amplification products from each RNA sample in biological replicates. Sequence aligning and comparison was done using ApE and SnapGene Viewer (25, 50).

### SDS PAGE and Western Blotting Analysis

Whole-cell lysates of gonococci were harvested by pelleting cultures at a standardized optical density and resuspending cells in 2x Laemmli solubilizing buffer (BioRad, cat number) before storage at −20°C. Lysates were thawed and mixed with βME to a final concentration of 5%. Samples were then briefly centrifuged and boiled for 2 min at 100°C. Protein samples were separated on a precast 4-to-20% gradient polyacrylamide gel (BioRad, cat number) before transfer to nitrocellulose. Prior to transfer, the SDS PAGE electrophoresis gel was analyzed by stain-free method to ensure equivalent protein loading and separation. The blots were further stained with Ponceau S after transfer, to verify equal protein sample loading. To detect TdfJ, TbpB and Ngo1049, the blots were blocked in 5% (wt/vol) Bovine Serum Albumin (BSA) dissolved in 1x High Salt-Tris Buffer Saline with 0.1% Tween 20 (HS-TBST). The blots were probed with peptide- or protein-specific polyclonal antisera diluted in blocker or for 1 h at room temperature. Polyclonal Rabbit anti-TdfH and -TbpB were diluted at 1:2500, Guinea Pig anti-TdfJ was diluted at 1:200, Guinea Pig anti-Ngo1049 was diluted at 1:500 and Guinea Pig anti ZnuA was diluted at 1:1000 in a 5% BSA blocker. The blots were washed three times with 1x HS-TBST and then probed with AP-conjugated anti-host (guinea pig or rabbit) IgG secondary antibodies at 1:5000 in BSA blocker for 1 h at room temperature. The blots were washed again after secondary antibody incubation and developed using either nitroblue tetrazolium (NBT)–5-bromo-4-chloro-3-indolylphosphate (BCIP) for AP-conjugated secondary antibodies or the ECL substrate (BioRad) for the HRP labelled secondary antibody after incubating for about 10 to 15 mins at room temperature with gentle shaking. To detect recombinant His-NgFur, the blots were blocked in 5% (wt/vol) skim milk in 1x HS-TBST and probed with anti-His HRP antibody diluted at 1:1000 in blocker for 1 hour at room temperature. The NgFur blots were developed using ECL substrate from BioRad. Blots were imaged on a Bio-Rad ChemiDoc gel imaging system using the Colorimetric or auto-ECL detection.

### Coomassie Stain

To detect purified protein His-NgFur, which does not contain tryptophan residues, SDS-PAGE gels of purification products of NgFur, were stained with Coomassie blue (0.25% Coomassie R-250, 50% methanol, 10% glacial acetic acid). The stain free gel imaging system on the Chemidoc gel imager only works for proteins that are rich in tryptophan and since NgFur has no tryptophan we resorted to using the Coomassie staining of purified His-NgFur (51). The gel was stained for 1 hour at room temperature and de-stained in 20% methanol and 5% acetic acid overnight at 4°C to minimize background. The stained gels were then imaged under the colorimetric setting in a Chemidoc gel analyzer to look for purified Fur.

### Expression and Purification of Recombinant His-NgFur

The coding sequence for FA1090 fur was PCR amplified with primers oGSU261 and oGSU262 (Table 2). The resulting PCR product was digested with EcoR1 and BamH1 and cloned into the pET19b vector that contains a 10X polyhistidine tag at the N-terminus, to result in the plasmid pGSU606 (Table 1). This plasmid after sequence confirmation was purified and transformed into BL21(DE3 pLysE) competent cells. The His-NgFur protein was expressed in a starter culture of the BL21 (pGSU606) plasmid in 5 mL Luria Bertani broth supplemented with 100 µg/mL of carbenicillin; this starter culture was sub-cultured into Terrific Broth (TB) supplemented with 100 µg/mL carbenicillin. Expression culture was grown with 1 mM inducer-isopropyl β-d-1 thiogalactopyranoside (IPTG) at 37°C for 4 hours. Cell pellets were harvested by centrifugation and stored at -80°C before lysis. Cell pellets were re-suspended in lysis buffer (20 mM HEPES, 250 mM NaCl, pH 8.0, 1 mM phenylmethylsulfonyl fluoride (PMSF), 1 U/µL Pierce Universal Nuclease for Cell lysis) and lysed using a Dounce homogenizer, with 10 mL of buffer used per gram of cell pellet (35). The resulting cell suspension was mechanically lysed by sonication at 50% amplitude at Pulse-time mode, with sonication at 30 second intervals for 10 minutes. Insoluble fraction was removed by centrifugation at 12,500xg for 1 hour at 4°C and the resulting clear supernatant was mixed with pre-equilibrated nickel-nitrilotriacetic acid (Ni^2+^-NTA) resin, 10 mM 2-Mercaptoethanol (βME) and 1 mM PMSF at 4°C overnight. The Ni^2+^-NTA resin was collected in a chromatography column and washed with 30 column volumes (cv) and eluted with 10 cv total with wash buffer (20 mM HEPES, 150 mM NaCl, pH 8.0, 10 mM βME, 1 mM PMSF) containing 50 to 500 mM imidazole. Clean His-tagged NgFur eluted with buffer containing 200-, 300- and 400-mM imidazole. These fractions were pooled together and dialyzed against 1x Phosphate Buffer Saline (pH 8.0), overnight at 4°C to remove the imidazole. The purity of purified His-NgFur was determined by SDS-PAGE electrophoresis followed by staining with Coomassie blue. No reducers such as, Dithiothreitol (DTT) and βME were added to the dialyzed His-NgFur as they would interfere with downstream assays. Protein concentration was measured and His-NgFur was aliquoted, and flash-frozen at 80°C.

### DNA-Protein Interaction (DPI) ELISA

A biotinylated DNA probe was produced by PCR amplification of the promoter regions of *tbpB*, *ngo1049*, *tdfJ* and *rmpM*, using 5’ BIOTIN-TEG tagged reverse primers, oGSU300 (P_tdfJ_), oGSU302 (P_ngo1049_), oGSU304 (P_tbpB_) and oGSU386 (P_rmpM_). These primers were designed and validated on IDT with HPLC purification prior to ordering. PCR amplified biotinylated DNA probes were bound to a streptavidin-coated 96-well plate (Pierce High-Capacity Plates, Cat: 15500) and ELISA was carried out according to the protocol by Brand, L. H., et al (30) with the following modifications. Biotinylated PCR DNA, diluted to 12.5 pmols in 1x Low-Salt Tris Buffer Saline (LS-TBS) was bound (100 µL per well) to a pre-washed streptavidin-coated high-capacity plate (∼125 pmol BIOTIN binding capacity) and incubated for 2 hours at room temperature with gentle shaking. After 2 hours the plate was washed in a programmed plate washer 5 times at 300 µl each with 1x phosphate buffer saline (PBS) pH 7.2. A 5 % milk in 1x LS-TBS with 0.05 % Tween 20 blocker was added at 200 µL per well and incubated at 4°C overnight. His-NgFur at the highest required concentration (20µM) was loaded with Mn^2+^ from MnCl_2_ in the molar ratio of 1:2 Fur:Mn^2+^ (40µM MnCl2) and left in a nutator at 4°C for 1 hour. Mn^2+^ loaded His-NgFur was then serially diluted in Fur dialysis buffer, 1xPBS at pH=8.0 to the desired gradient concentrations ranging from 20µM to 10µM Fur and added to the DNA bound plate for 1 hour at room temperature with gentle shaking. The plate was washed five times with 1X PBS supplemented with 0.05% Tween 20 (PBS-T) in the plate washer. An anti-His HRP labelled antibody (BioLegend Cat: J099B12) was diluted in blocker at 1:1000 and used to detect the DNA bound His-NgFur, by incubating for 1 hour at room temperature with gentle shaking. The plate was washed after the anti-His antibody step in 1x PBS-T and photometric detection was performed with the Slow TMB substrate (Thermo Fischer Cat: 34024). The reaction was stopped with 0.18 M sulfuric acid and the resulting absorbance was read at 450 nm in a plate reader. The binding affinity K_D_ was determined through competition of biotinylated P_tbpB_, P_ngo1049_, P_tdfJ_ and P_rmpM_ with their respective unlabeled 10x excess PCR promoter DNA counterparts. The binding curves were plotted and His-NgFur K_D_ to P_tdfJ_ was determined by a Linear regression model curve fitting with the Hill equation plot in GraphPad PRISM (52). The specificity of His-NgFur to the promoters were determined through binding to biotinylated scrambled sequence of P_tdfJ_ Fur and Zur boxes and P_tbpB_ Fur box. The binding curves were generated by determining the µg of protein bound to an ELISA plate through a standard curve plot for His-NgFur in biological triplicates and technical quadruplets. The K_D_ for the promoters were also done in biological triplicates and technical duplicates to obtain statistical significance.

### Fur Scrambled Promoter ELISA

For creating scrambled versions of P_tdfJ_ and P_tbpB_ promoter Fur and or Zur binding regions, we synthesized gene blocks through IDT. Gene blocks gbGSU015 to gbGSU019 and gbGSU021 containing modified Fur and Zur box regions on P_tdfJ_ to varying capacity with sequence scarmbling on gbGSU015 being the least. gbGSU021 was made where 100% scrambling of Fur and Zur box regions of P_tdfJ_ were achieved. Scrambling was done by modifying the consensus repeat regions for Fur or Zur box, that is AT strings to G’s and C’s. A similar approach was taken to synthesize gene blocks gbGSU014 and gbGSU020 for scrambled Fur box of P_tbpB_, where gbGSU020 is the most scrambled P_tbpB_ Fur box. These gene blocks were prepared according to manufacturer’s protocol (IDT) and then TOPO cloned using the Invitrogen Zero BLUNT TOPO cloning vector (Cat: 450245). Top 10 or DH5-α E. coli were transformed with the TOPO cloned DNA and selected on LB plates with Kanamycin at 50µg/mL overnight at 37°C. The transformed colonies were confirmed through sequencing and resulted in plasmids pGSU623 to pGSU629 described in Table 1. pGSU623 and pGSU628 contain plasmid DNA with scrambled P_tbpB_ and plasmids pGSU624 to pGSU67 and pGSU629 contain plasmid DNA with versions of scrambled P_tdfJ_. Plasmid DNA from these plasmids were isolated by growing them overnight in liquid LB media at 37°C and using the Zymoresearch Plasmid Mini Prep Kit. Plasmid DNA were them amplified by PCR using labelled Biotinylated primers for DPI-ELISA (oGSU299-oGSU300 for P_tdfJ_ and oGSU303 -0gUS304 for P_tbpB_) using the NEB Phusion Polymerase (Cat: M0530S) and 10mM dNTP. This resulted in biotinylated scrambled versions of P_tbpB_ and P_tdfJ_ ready to be bound on the streptavidin ELISA Plates for a Fur DPI-ELISA as described in the previous section (Section-11). DPI ELISA was used to measure the absorbance values for Fur binding and resulting values were plotted using Excel as bar graphs.

### In Silico Promoter Analysis and Prediction

Online promoter prediction software such as BPROM (22) and PRODORIC-Virtual Footprint (23) were used to identify the putative promoter elements and DNA Fur- and Zur-binding sites, along with consensus sequences, through pattern recognition. In addition to online prediction tools, predicted promoters from RNA Sequencing were analyzed using Geneious Prime, WebLogo 3 and MEME Suite (21, 53–55), to generate a consensus logo for Ngo FA1090 Fur and Zur box using alignment to known sequences of consensus.

### Statistical Analysis

Mean with standard deviation (SD) was calculated and statistical significance was analyzed by Student’s t tests or one-way ANOVA. Significance was set at p < 0.05. Prism software version 3.0 was used for graphing gene expression bar graphs and protein binding affinity K_D_ calculations

